# Exercise-stimulated Resolvin Biosynthesis in Adipose Tissue is Abrogated by High Fat Diet-induced Adrenergic Deficiency

**DOI:** 10.1101/2024.08.14.608014

**Authors:** Ernesto Pena Calderin, Jing-Juan Zheng, Nolan L. Boyd, Will Lynch, Brian Sansbury, Matthew Spite, Bradford G. Hill, Jason Hellmann

**Author notes:** To whom correspondence should be addressed: Jason Hellmann, Ph.D., 580 S. Preston St. Rm 204F Delia Baxter II Building University of Louisville Louisville, KY 40202.

## Abstract

**Objective:** Diet-induced white adipose tissue inflammation is associated with insulin resistance and metabolic perturbations. Conversely, exercise (Exe) protects against the development of chronic inflammation and insulin resistance independent of changes in weight; however, the mechanisms remain largely unknown. We have recently shown that, through adrenergic stimulation of macrophages, exercise promotes resolution of acute peritoneal inflammation by enhancing the biosynthesis of specialized pro-resolving lipid mediators (SPMs). In this study, we sought to determine if exercise stimulates pro-resolving pathways in adipose tissue and whether this response is modified by diet. Specifically, we hypothesized that high fat diet feeding disrupts exercise-stimulated resolution by inhibiting adrenergic signaling, priming the development of chronic inflammation in adipose tissue (AT).

**Approach and Results:** To explore the dietary dependence of the pro-resolving effects of Exe, mice were fed either a control or high-fat diet (HFD) for 2 weeks prior to, and throughout, a 4 wk period of daily treadmill running. Glucose handling, body weight and composition, and exercise performance were evaluated at the end of the feeding and exercise interventions. Likewise, catecholamines and their biosynthetic enzymes were measured along with AT SPM biosynthesis and macrophage phenotype and abundance.

When compared with sedentary controls (Sed), macrophages isolated from mice exposed to 4 wk of exercise display elevated expression of the SPM biosynthetic enzyme *Alox15,* while whole AT SPM levels and anti-inflammatory CD301+ M2 macrophages increased. These changes were dependent upon diet as 6 wk of feeding with HFD abrogated the pro-resolving effect of exercise when compared with control diet-fed animals. Interestingly, exercise-induced epinephrine production was inhibited by HFD, which diminished expression of the epinephrine biosynthetic enzyme phenylethanolamine N-methyltransferase (PNMT) in adrenal glands.

**Conclusion:** Taken together, these results suggest that a diet high in fat diminishes the pro-resolving effects of exercise in adipose tissue via decreasing the biosynthesis of catecholamines.

**GRAPHICAL ABSTRACT:** 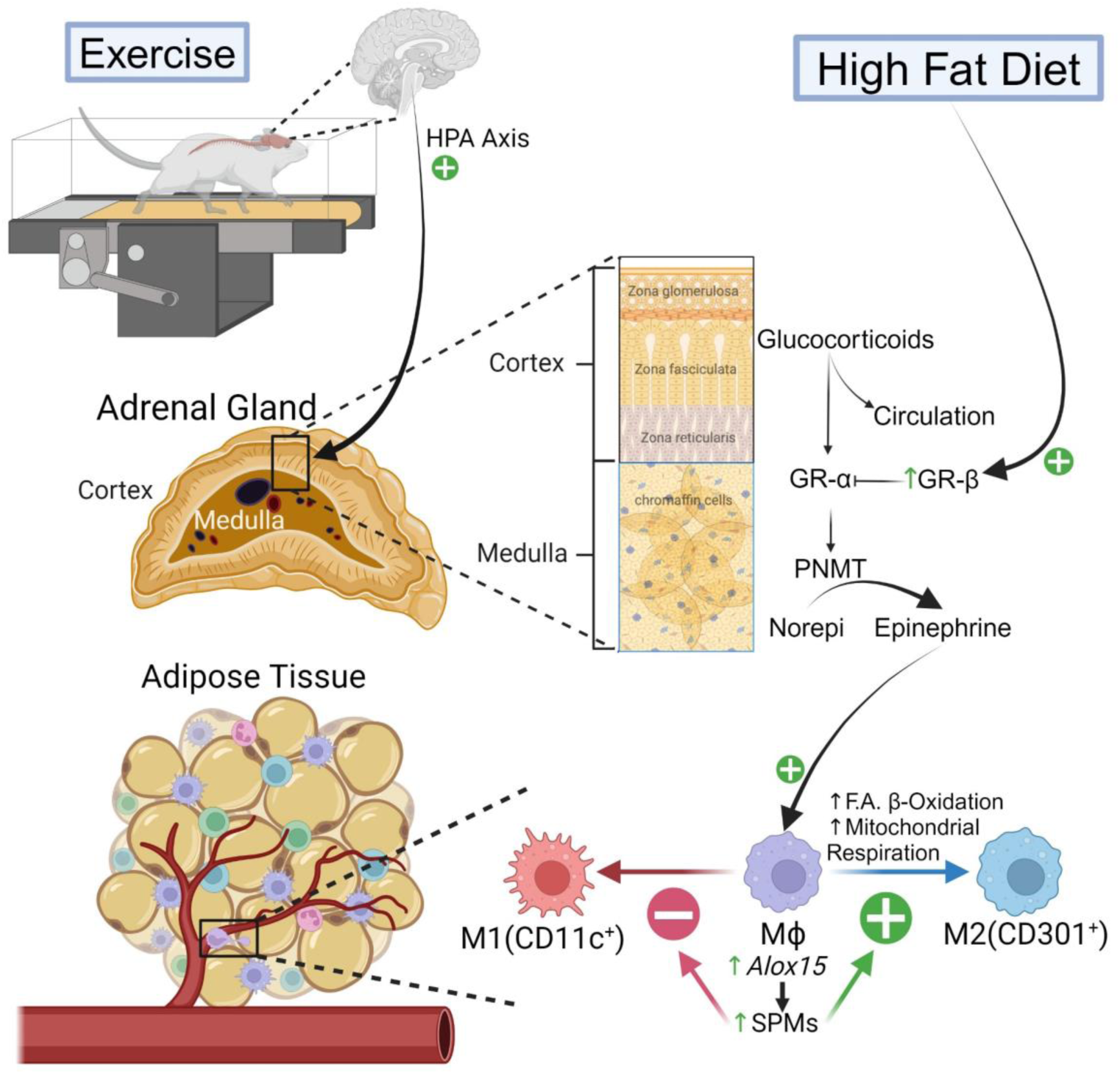

## 1. INTRODUCTION

According to data from the 2017-2018 National Health and Nutrition Examination Survey (NHANES) and the World Health Organization, the prevalence of obesity is a significant global health concern, impacting 42.4% adults in the United States and 13% worldwide. Obesity, defined as having a body max index equal to or greater than 30 kg/m^2, 1–3^ is on an alarming trajectory with projections indicating that nearly half of the US population may be obese by 2030^4^. Beyond its evident impact on weight, obesity is associated with a heightened risk for developing hypertension, heart failure, stroke, type 2 diabetes mellitus (T2D), non-alcoholic fatty liver disease, dyslipidemia, and certain types of cancers.^5^ These comorbidities place a tremendous burden on healthcare systems globally. The primary drivers behind the escalating rates of obesity and obesity-induced comorbidities are poor dietary habits and a sedentary lifestyle.^6^ Understanding mechanisms underlying the detrimental effects of a poor diet and sedentary behavior is crucial for preventing and managing obesity, promoting overall health, and alleviating healthcare strain.

In response to changes in nutritional status, the adipose tissue undergoes dynamic remodeling, which during chronic nutrient excess can become perturbed resulting in immune cell recruitment and the development of adipose tissue inflammation. Adipose tissue inflammation is well documented to contribute to the development of obesity-induced T2D and related comorbidities. In addition to immune cell recruitment, inflamed obese adipose tissue is also characterized by an imbalance in lipid mediator (LM) production with an increase in omega-6 polyunsaturated fatty acid (PUFA)-derived LMs and decreased production of omega-3 PUFA-derived specialized pro-resolving mediators (SPMs).^7,8^ This imbalance results in defective inflammation resolution programs, contributing to elevated M1 inflammatory macrophage abundance and the development of chronic inflammation, insulin resistance, and T2D. Several reports have shown that, in either genetically obese or diet-induced obese mice, restoring the LM imbalance to favor resolution with SPM treatment (e.g., lipoxins^9^, maresins,^10^ resolvins,^11,12^ or SPM precursors^13^) ameliorates adipose tissue inflammation with a reduction in pro-inflammatory cytokines (TNF-α, IL-6, MCP-1, and IL-1β), increased M2:M1 adipose tissue macrophage (ATM) ratio, and increased anti-inflammatory cytokine IL-10 production. Conversely, as adipocytes decrease in size during nutrient utilization there is a reduction in pro-inflammatory M1 macrophage abundance with an increase in the abundance of pro-resolving M2 macrophages. This results in a decrease in the release of pro-inflammatory cytokines TNF-α and IL-6 (along with a decrease in proinflammatory and insulin-desensitizing adipokines leptin and RBP4), and a concomitant increase in anti-inflammatory cytokines (i.e. IL-10 and IL-1RA) and insulin-sensitizing hormones such as adiponectin.^14–16^ These findings suggest a critical role of SPMs in mediating immune cell phenotype and, when absent, subsequent propensity for the development of adipose tissue inflammation during excess nutrient consumption.

Our laboratory has recently shown in a cross-sectional analysis of NHANES data that plasma levels of omega-3 PUFAs (SPM precursors) modify the inverse relationship between cardiorespiratory fitness and systemic inflammation, which suggest that the anti-inflammatory effects of exercise are mediated by omega-3 PUFAs.^17^ Additionally, several recent studies have found that circulating levels of SPMs (i.e., RvD1 and LXA_4_) and many of their precursors are elevated with exercise in healthy, exercise-trained and untrained human subjects.^18–21^ In preclinical mouse models of acute inflammation, we found that exercise enhances macrophage phagocytosis and hastens resolution through epinephrine-induced stimulation of SPM biosynthesis.^22^ Additionally we recently reported that SPMs promote mitochondrial respiration in macrophages, an emerging characteristic of M2 pro-resolving macrophages.^23^ Collectively, these findings suggest that exercise promotes macrophage SPM biosynthesis, mitochondrial respiration, and resolution of acute inflammation.

In this study, we aimed to assess the proresolving effects of exercise in adipose tissue and to identify their dependence on dietary composition. Importantly, to prevent the obfuscating effects of overt obesity, which is associated with increased inflammation, we chose to examine whether a relatively short duration of high fat diet influences resolution programs in adipose tissue. Our findings indicate that a diet high in fat diminishes the pro-resolving effects of exercise in adipose tissue via decreasing catecholamine biosynthesis.

## 2. MATERIALS AND METHODS

### 2.1 Animals

Male and female FvB/NJ (WT) mice were purchased at 6 wk of age from The Jackson Laboratory (strain no. 001800; Bar Harbor, ME, USA) and were housed with up to 3 littermates per cage in standard housing conditions – namely, in a temperature-controlled environment (21°C year-round), with a 12:12-h light/dark cycle, and with *ad libitum* access to food and water. For exercise time course experiments, 8-9 wk old male mice were maintained on a normal chow (NC) diet and throughout 1, 2, or 4 wk of treadmill exercise. To test dietary dependence where synthetic diets and exercise training were administered, standard, normal chow (NC) diet (12.73% kcal; cat. no. 5010; Lab Diets, Richmond, IN, USA) was used from weaning age till 8 weeks of age, when male mice were randomly assigned to either a 10% kcal low fat diet (LFD) (cat. no. D12450B; Research Diets, New Brunswick, NJ, USA) or a 60% kcal high fat diet (HFD) (cat. no. D12492; Research Diets) for 2 wk prior to, and throughout 4 consecutive wk of either treadmill exercise or sedentarism. In a separate experiment, 8-wk old male (experimental results presented in **Fig. 5**) and female mice (experimental results presented in **Fig. S5-E** in the Supplemental Manuscript) were fed HFD for 6 wk before, and throughout a daily (once a day, Monday through Friday), week-long administration of either sterile normal saline (vehicle control) or epinephrine (0.1 mg/kg body weight, i.p.). No injections were administered over the weekend after the 5 daily doses, and then 1 last dose was administered on Monday, followed by euthanasia 24 h later. This same experimental setup was repeated on a separate cohort where only male mice were administered either vehicle (10% v/v Kolliphor HS 15 (cat. no. 42966-1KG; Sigm-Aldrich, St. Louis, MO, USA), 10% v/v Kolliphor EL (cat. no. C5135-500G; Sigma-Aldrich), 20% v/v Kollisolv PEG400 (cat. no. 06855-1KG; Sigma-Aldrich), 6% v/v DMSO (cat. no. D12345; Invitrogen, Waltham, MA, USA), 54% v/v sterile normal saline) or 50 mg/kg ML351 (cat. no. HY-111310 (i.p) in the morning (11 AM), followed by a 0.1 mg/kg epinephrine (cat. no. E4642-5G; Sigma-Aldrich) injection 2 h later. In separate experiments where either peritoneal macrophages (PM) or stromal vascular fraction (SVF) cells of the epididymal white adipose tissue (eWAT) were extracted, male FvB/NJ mice 11-12 and 25 wk old, respectively, were maintained on a NC diet at standard housing conditions until euthanasia. All animal procedures were approved by the University of Louisville Institutional Animal Care and Use Committee (IACUC).

### 2.2 Mouse treadmill Exercise, GTT, ITT, fasting blood glucose, DEXA scan, and hematological analysis

Mice were subjected to treadmill exercise as previously described ^22^. After 2 d of familiarization with the treadmill environment (Exer 3/6 Treadmill; Columbus Instruments, Columbus, OH, USA) a graded, maximal exercise capacity test (ECT) was performed to assess baseline exercise capacity (as measured by distance run and work performed till exhaustion) in all mice. Then, 48 h following the baseline ECT, mice were subjected to either daily (Mon-Fri) treadmill exercise at 75% of their average maximal ECT speed or were left sedentary for the indicated number of weeks. After the exercise training period, a final ECT was administered to assess changes in exercise performance. Immediately within 1 minute after exhaustion, blood lactate levels were measured in venous blood (tail vein) using a hand-held Lactate Plus lactate meter (Nova Biomedical, Waltham, MA). See reference^22^ for familiarization, ECT and daily running protocols, respectively.

A glucose tolerance test (GTT) and an insulin tolerance test (ITT) were performed on the third week of exercise training (1 week before euthanasia) with 72 h elapsing between each test. Mice were acclimated to the testing environment 16 h prior. For the GTT, mice were fasted for 6 h before measuring baseline fasting blood glucose levels and then administered an intraperitoneal bolus of 1 mg D-glucose (cat. no. G7528-250G; Sigma-Aldrich) per gram of body weight in sterile normal saline with subsequent measuring of blood glucose levels with a hand-held glucometer (Aviva Accu-Chek; Roche, Basel, Switzerland) at the indicated time points. For the ITT, non-fasted mice were given 1.5 U/kg body weight Humulin™ R (cat. no. R-100, Eli Lilly, IN, USA) in sterile normal saline intraperitoneally and blood glucose levels were measured at the indicated time points. Dual-energy x-ray absorptiometry (DEXA) scans (PIXImus, GE Medical Systems, Chicago, IL, USA) to determine percent lean and fat mass were taken by briefly anesthetizing mice with 2% isoflurane on the third day of the third week of exercise training. All other endpoint analyses were performed in mice 24 h after the last exercise bout.

Fresh mixed venous blood was collected following euthanasia in unfasted animals via intracardial puncture (into right ventricle) using syringes with 1” 25-G needles, and collected into tubes containing 1.5 – 2.2 mg of EDTA per mL of whole blood (3.2 – 5 mM final [EDTA]) to prevent coagulation. Plasma was separated by centrifugation of non-coagulated whole blood at 1500 RCF for 15 min at 4 °C. Following isolation, plasma was used in measuring circulating insulin, adiponectin, creatinine, free fatty acids (FFA) and glycerol. A separate aliquot of uncoagulated fresh whole blood was used to measure monocytes in a Hemavet 950 (Drew Scientific, Miami Lakes, FL, USA)

### 2.3 Stromal vascular fraction (SVF) and adipocyte fraction extraction from epididymal white adipose tissue (eWAT)

Epididymal white adipose tissue (eWAT) was dissected under aseptic conditions, and 100-200 mg of tissue was finely minced on a borosilicate glass dish surface with 1 mL of ice-cold digestion buffer composed of DPBS containing Ca^2+^ and Mg^2+^ (cat. no. 14040133, Thermo Fisher, Waltham, MA, USA) and 1% v/v heat-inactivated fetal bovine serum (FBS) (cat. no. S11150H, R&D Systems, Minneapolis, MN, USA). Tissue homogenates were kept on ice for no more than 1 h until all samples were simultaneously digested at 37°C for 30 min, while gently inverting and swirling tubes every 5 min. Digestion solution contained a ratio of 2 mg (0.5 – 5.0 FALGPA units) of collagenase (*C. histolyticum* extract; cat. no. C6885-25MG, Millipore Sigma, Burlington, MA, USA) per 100-200 mg of eWAT, at a collagenase concentration of 1 mg/mL in digestion buffer. The digestion reaction was stopped by adding EDTA (cat. no. 324506-100ML; EMD Millipore, Burlington, MA, USA) to a final concentration of 10 mM and by immediately placing samples on ice for 5 min. Cell suspensions were passed through a 100 μm cell strainer to eliminate undigested tissue and subsequently centrifuged for 5 min at 680 RCF at 4°C. All samples were kept at 4°C for the reminder of the experiment. The supernatant was carefully aspirated and collected to separate out floating adipocytes, which were collected for further analysis. Cell pellets containing the stromal vascular fraction (SVF) were washed by resuspension in 5 mL post digestion buffer (composed of Ca^2+^ and Mg^2+^ free DPBS supplemented with 1% v/v FBS), followed by centrifugation and removal of the supernatant. Last, the SVF cell pellets were resuspended in 1 mL and cells were counted using an automated cell counter (Bio-Rad TC-10).

### 2.4 F4/80+ adipose tissue macrophage (ATM) purification from eWAT stromal vascular fraction

ATM isolation was performed using EasySep^TM^ PE Positive Selection Kit (cat. no. 17666, STEMCELL Technologies, Vancouver, BC, Canada) according to manufacturer recommendations. All steps were conducted at room temperature and centrifugation speed and time were kept constant at 680 RCF and 5 min, respectively. Briefly, 1E6-9E6 SVF cells were loaded onto 17×100mm polystyrene round-bottom test tubes and FcR receptors blocked by incubation with 1.5 μg of a rat anti-mouse CD16/CD32 monoclonal antibody (cat. no. 14-0161-82, Thermo Fisher) for 10 min at room temperature in 250 μL post digestion buffer. Next, 0.6 μg of a rat anti-mouse F4/80-PE antibody was incubated for 15 min at room temperature and samples were washed with 3 mL of EasySep^TM^ buffer (ESB) (cat. no. 20144, STEMCELL Technologies), centrifuged, supernatant discarded, and cell pellet (1.2E7 to 1.7E7 cells) resuspended in 225 μl ESB. F4/80^+^ ATM were selected by incubating 25 μl of PE selection cocktail (EasySep^TM^ PE Positive Selection Kit) for 15 min, followed by a 10 min incubation with

24.38 μL EasySep Dextran RapidSpheres. Sample volume was brought to 5 mL and test tubes were placed in a magnetic block for 10 min (“The Big Easy” EasySep magnet, cat. no. 18001, STEMCELL Technologies). Tubes were gently rocked to allow F4/80^+^ ATM to be magnetically pulled towards the wall of the test tubes. After incubation, the supernatant (containing F4/80^-^ cells) was discarded and remains cells were washed with 5 mL ESB by resuspending and pulling them towards the wall of the test tubes once again. This wash step was repeated 2 more times to increase F4/80^+^ selection enrichment. At the end of the last wash, cells were resuspended and enumerated.

Loading of 1E6-9E6 SVF cells (digested from 200-400 mg of tissue) typically yielded 0.5E6-1E6 ATM cells.

### 2.5 Flow Cytometric Analysis of eWAT SVF or isolated ATM

After their respective isolations and purifications, 0.25E6 – 1E6 SVF or ATM cells were resuspended and maintained in ice-cold FACS buffer (Ca^2+^ and Mg^2+^ free DPBS supplemented with 1% v/v FBS) for the remainder of the flow cytometric analysis protocol. Samples were then centrifuged at 680 RCF for 4 min at 4°C. After removing the supernatant, cells were FcR-blocked for 10 min by incubating 0.5 μg of a rat anti-mouse CD16/CD32 monoclonal antibodies in 50 μl. After FcR blocking, 50 μl of a staining solution containing the following primary flow antibodies – F4/80-PE, CD11c-PE/Cy7 and CD301-APC – was added for a total staining volume of a 100 μl, with each antibody at a final concentration of 20 μg/ml. See Table S4 in Supplemental Methods for detailed antibody information. Unstained and single-stain control samples were prepared alongside in order to set positive and negative fluorescence gates and to calculate spectral overlap. After 30 min incubation, samples were washed with 5 volumes of FACS buffer, and cell pellets were resuspended in 300 μl of FACS buffer. Samples were submitted for flow cytometric analysis using a BD LSRFortessa X-20 flow cytometer (BD Bioscience, Franklin Lakes, NJ, USA) with BD FACSDiva v8.0.3 acquisition software. Data were analyzed using BD FACSDiva BD v8.0.3 and FlowJo v10.8.1.

### 2.6 Lipid mediator quantification by targeted liquid chromatography-tandem mass spectrometry (LC-MS/MS)

To quantify differences in lipid mediators, mice were euthanized and epididymal white adipose tissue (eWAT) was immediately snap-frozen and stored at -80°C until subjected to LC-MS/MS analysis. Lipid mediator quantification was performed by 2 independent laboratories utilizing similar methodological approaches and instrumentation. On the day of extraction, 400 µL of ice-cold HPLC-grade MeOH was added to 50–100mg of tissue and minced and homogenized. Protein was precipitated at -20°C for 1 h after the addition of 600 µL of MeOH containing 500 pg of deuterium-labeled internal standards (i.e. Resolvin D_2_-d_5_, Resolvin D_3_-d_5_, Maresin 1-d_5_, Maresin 2-d_5_, Lipoxin A_4_-d_5_, Resolvin E_1_-d_4_, 5(S)-HETE-d_8_, 15(S)-HETE-d_8_, (+)11(12)-EET-d_11_, 15-deoxy-Δ12,14-Prostaglandin J_2_-d_4_, Prostaglandin E_2_-d_4_, and Leukotriene B_4_-d_4_) to calculate extraction efficiency. Protein precipitates were pelleted by centrifuging samples at 6200 RCF for 10 min at 4°C and supernatants were transferred into 12 mL borosilicate glass round-bottom tubes. Samples were then acidified by the addition of pH 3.5 HPLC-grade H_2_O and promptly loaded onto Biotage ISOLUTE C18 SPE columns (cat. no. 220-0020-c, Biotage, Uppsala, Sweden), which had been pre-equilibrated with 3 mL of HPLC-grade MeOH and 3 mL HPLC-grade H_2_O. After samples were loaded, columns were then neutralized with 5 mL of HPLC-grade H_2_O and subsequently treated with 5 mL of HPLC-grade hexane to remove neutral lipids. Lipid mediators of interests were then eluted and collected by the addition of 5 mL HPLC-grade methyl formate. Samples were dried to completion with a gentle stream of N_2_ gas using a Biotage TurboVap Classic and immediately resuspended in 50 μL of HPLC-grade MeOH:H_2_O (50:50 v/v). For LC-MS/MS analysis, samples were injected onto a Kinetex Polar C18 HPLC column (100 mm length x 3 mm diameter; 2.6 μm particle size; 100 Å pore size) (cat. no. 00D-4759-Y0, Phenomenex, Torrance, CA, USA) maintained at 59.0°C using a Shimadzu LC-20AD with a SIL-20AC autoinjector (Shimadzu, Kyoto, JP), coupled to a 5500 Qtrap (Sciex, Toronto, ON, Canada). Targeted LC-MS/MS analysis was carried first by gradient elution of lipid mediators with a constant 0.5 mL/min mobile phase delivery, initially consisting of 45:55:0.01 (v/v/v) methanol:water:acetic acid, which was then ramped up to 80:20:0.01 over 16.5 min, and then to 98:2:0.01 over the next 2 min. The 5500 Qtrap was operated in negative polarity mode. All data were acquired using Analyst v1.7.1 and analyzed with Sciex OS-Q v3.0.0.3339 software (Sciex).

Each analyte was identified using multiple reaction monitoring (MRM) and enhanced product ion (EPI) mode (Supplemental Tables S1-S3) with retention time (RT) matching to synthetic standards (Fig. 2F). A given chromatographic peak was quantified if the following criteria were met: (1) RT was within ± 0.1 min of the synthetic standard’s RT; (2) chromatographic peak was composed of at least 5 points across baseline; (3) MS/MS spectra at the appropriate RT contained at least 6 diagnostic ions; (4) signal-to-noise ratio (SNR) was greater than 3; (5) the calculated analyte concentration was greater than the empirically-determined LLOD and LLOQ for the respective analyte. Sciex OS-Q Autopeak Integration Algorithm was used to integrate peaks and to generate peak area and SNR values.

Determination of the lower limit of detection (LLOD) and lower limit of quantitation (LLOQ) was carried out as recommended by the International Council for Harmonization analytical guideline Q2 (R2) (https://www.ich.org/page/quality-guidelines). Briefly, for all analytes, calibration standards (Cayman Chemical Company, Ann Arbor, MI, USA) were prepared by mixing twelve synthetic standards with a beginning concentration of 100 pg/μl (Standard 1) and serially diluted in HPLC-grade methanol (Sigma-Aldrich) by 1:2 dilution to a final concentration of 0.048 pg/μl (Standard 12). For each standard, an injection volume of 5 μl was used. At least 5 independent injections of each standard were used to generate data to determine LLOD and LLOQ for each analyte. From the accumulated data, the slope (S) of the calibration curve and the standard deviation (σ) of the y-intercept, which represents the blank condition, were calculated. LLOD and LLOQ were calculated using 3.3*σ/S and 10*σ/S, respectively.

An MS/MS spectra matching library was generated using Sciex OS LibraryView v1.4 (Sciex). To generate this library, 5 μl of synthetic standards at concentrations of 100pg/μl, 6.25pg/μl and 0.195pg/μl were injected to create a range of concentration-based MS/MS reference spectra that were added to the LibraryView database. This lipid spectra library was selected as part of the Sciex OS Process Method Library Search function. The search utilized the Smart Confirmation Search and Fit sorting options. The algorithm parameters Precursor Mass Tolerance and fragment Mass Tolerance were both set to ± 0.7Da. The Use Polarity Intensity Threshold option was set to 0.03. For each chromatographic peak quantified, a library matching score was generated and used to confirm the proper identification of the analyte based on a reference MS/MS spectra.

To determine pg of analyte in each sample, first, a 12-point calibration curve was created for each analyte in the standard mix by using a ratiometric method. With this method, a fixed (500) pg amount of internal deuterated standard (IS) was mixed with decreasing amounts of each synthetic standard, namely – 500, 250, 125, 62.5, 31.25, 15.63, 7.81, 3.91, 1.95, 0.98, 0.49, or 0.24 pg. The ratio of the peak areas was calculated and used as the Y-axis value, while the ratio of known pg of IS to synthetic standard on each calibration curve sample was used as the X-axis value to generate a calibration curve. All biological samples were spiked with the same pg of deuterated internal standards before extraction, and the peak area ratios of IS and analytes were calculated, and subsequently fitted into the calibration curve using a weighted (1/x) linear regression method to obtain pg of analyte in the biological sample. Each pg of analyte value was normalized to tissue mass processed.

For the generation of partial least-square determinant (PLSDA) 2D scores plot and heat maps in Fig. 1B and Fig 2H-I, normalized analyte concentrations from lipidomics datasets were uploaded to MetaboAnalyst 5.0 (www.metaboanalyst.ca). Analytes missing data in more than 50% of the samples were removed and missing values were imputed with 1/5 of the minimum positive value. Data were log transformed and autoscaled before analysis, as described in Fredman et al.^24^ Global changes between groups were compared using Student’s *t*-tests (P<0.05) and a fold-change threshold of 1.5.

**Figure 1.**
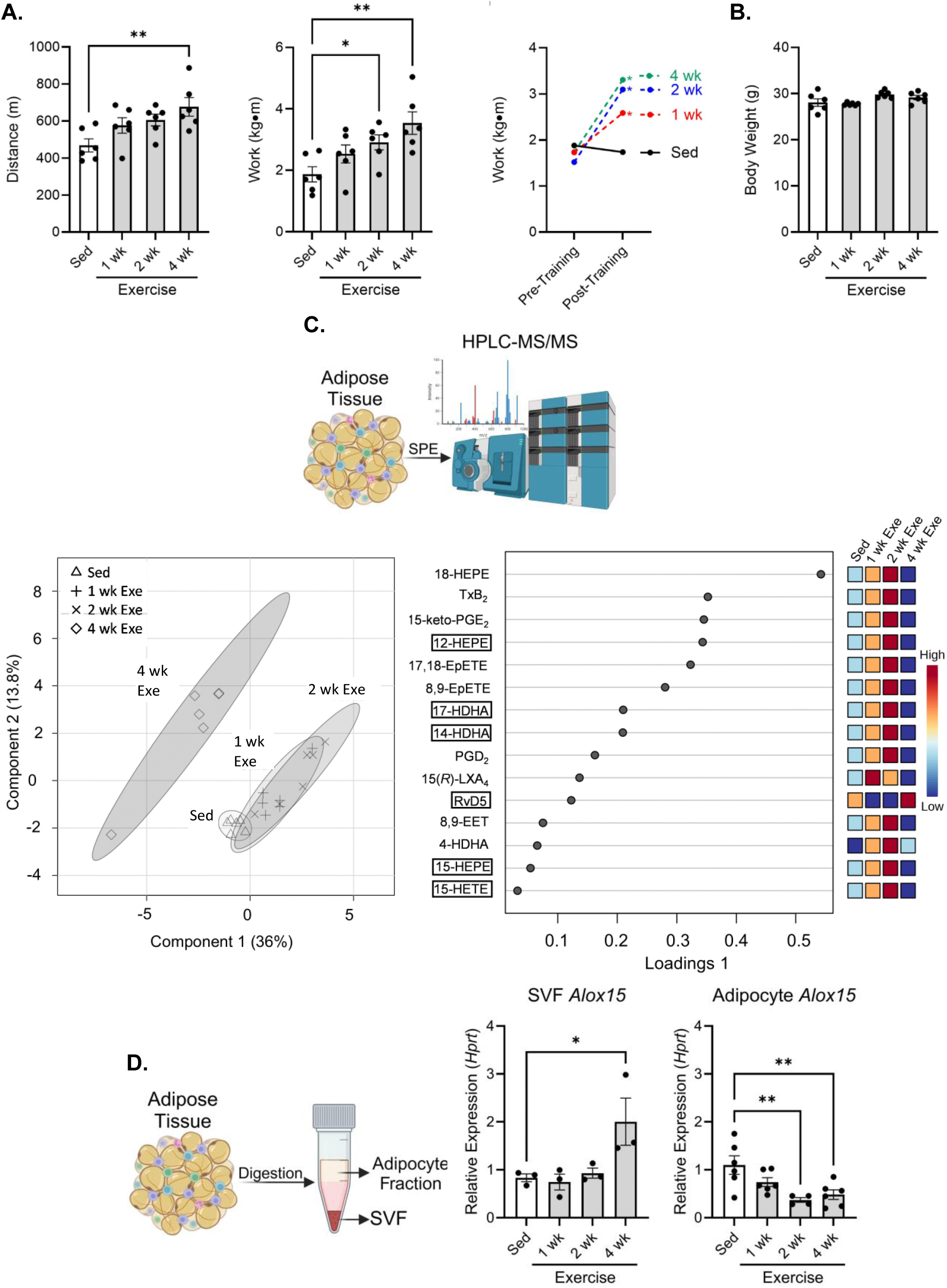
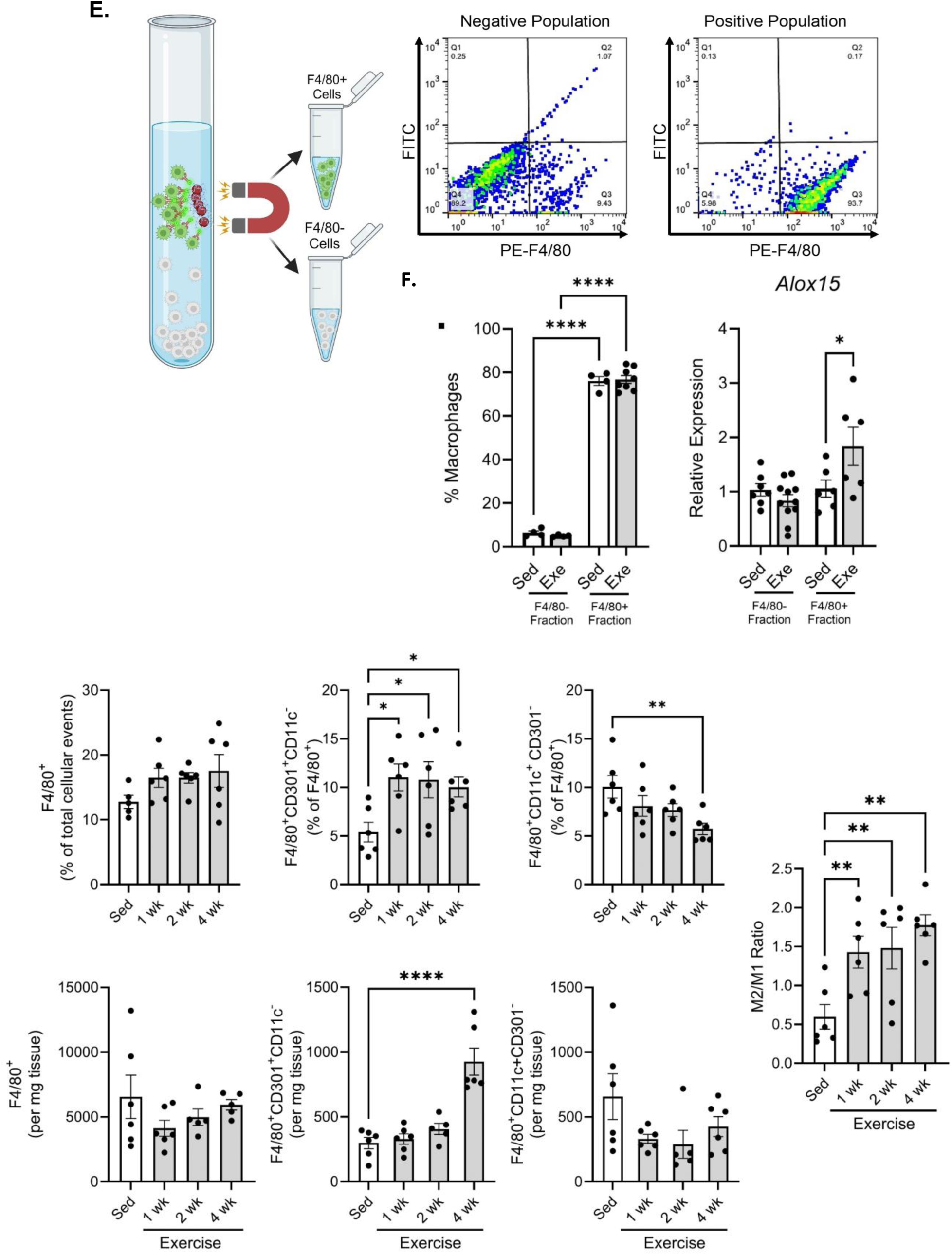
Exercise training decreases M1-polarized and increases M2-polarized adipose tissue macrophage content along with *Alox15* expression. (**A**) Distance and work performed during a graded, maximal exercise capacity test administered before and after the respective exercise training period, which lasted either 0 (Sed group), 1, 2, or 4 weeks (Exercise groups) in male mice. (**B**) Body weight measured at the end of the respective training periods. (**C**) Brief diagram outlining the main techniques used to generate the lipidomics data in this figure panel. 2D scores plot of sparse Partial Least Squares-Discriminant Analysis (sPLS-DA) and Loadings Plot of lipid mediator metabolipidomics from the eWAT. (**D**) Brief diagram outlining the main techniques used to generate the data in this figure panel – namely eWAT digestion and separation of the adipocyte and stromal vascular fractions, respectively, used to quantify *Alox15* mRNA at the end of the respective training period. (**E**) F4/80^+^ ATMs were extracted from the eWAT SVF of 4-weeks exercised male mice by magnetic beads and F4/80 expression levels from a representative sample are shown in a 2-D scatterplot for the F4/80^-^ and F4/80^+^ populations extracted. (**F**) The %F4/80^+^ ATM was quantified in the non-enriched and enriched fractions of sedentary and 4-weeks exercised male mice to verify ATM extraction efficiency. Subsequently, *Alox15* mRNA was quantified in the F4/80^-^ and F4/80^+^ SVF fractions of sedentary and 4-weeks exercised male mice. (**G.**) Brief diagram outlining the main techniques used to generate the data in this figure panel – namely eWAT digestion and SVF separation for flow cytometric analysis used to identify and quantify (as % of the respective parent population, and as cells per mg of eWAT digested) F4/80^+^ M1(CD11c^+^) or M2(CD301^+^) ATMs within the eWAT after the respective training period. Data expressed as mean ± SEM, *n* = 3-6 (A-D, F), *n* = 4-11 (E). \**P*<0.05, \*\**P*<0.01, \*\*\**P*<0.001, \*\*\*\**P*<0.0001; One-way (ANOVA with Holm-Šídák post-test).

**Figure 2.**
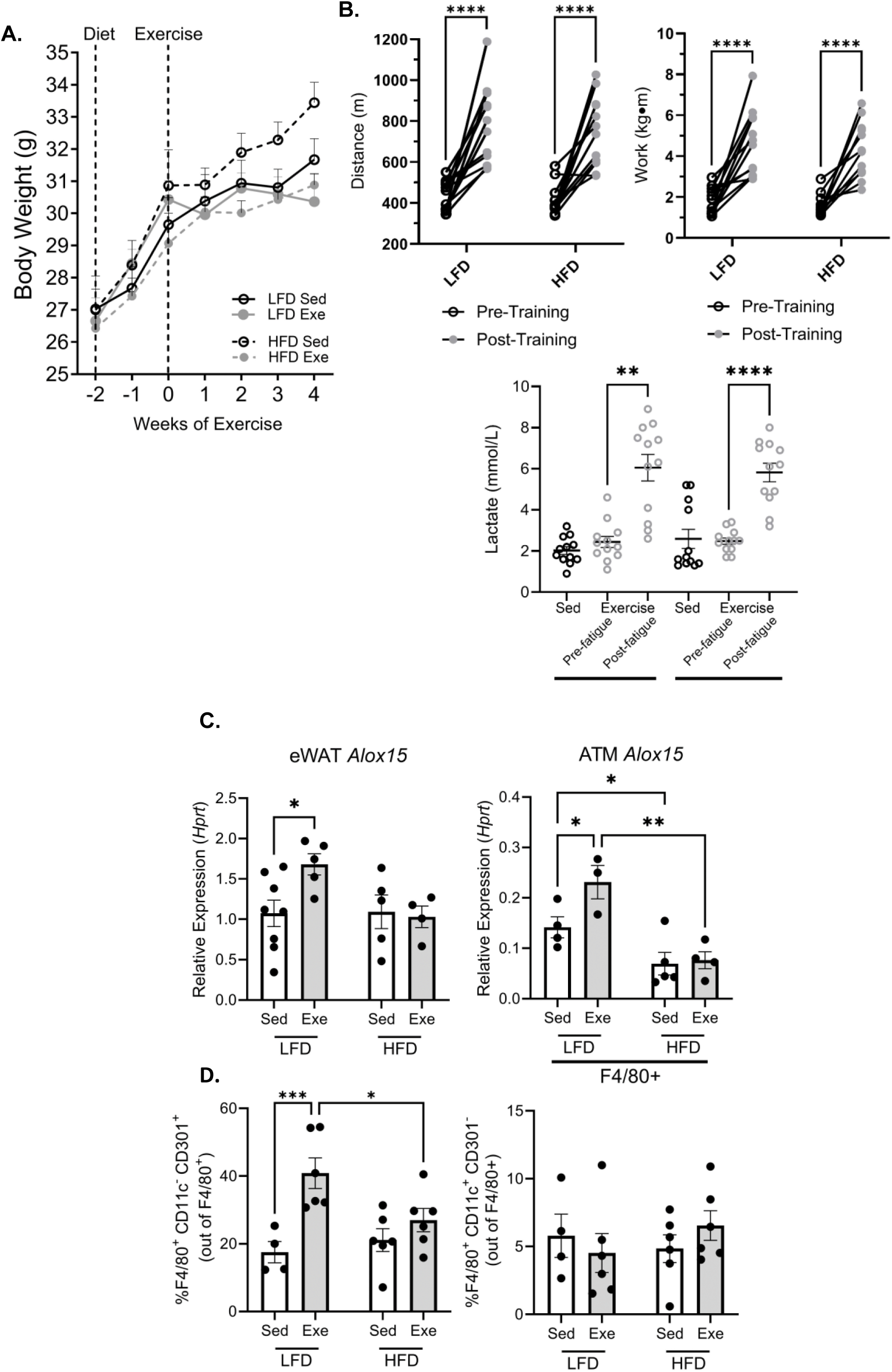
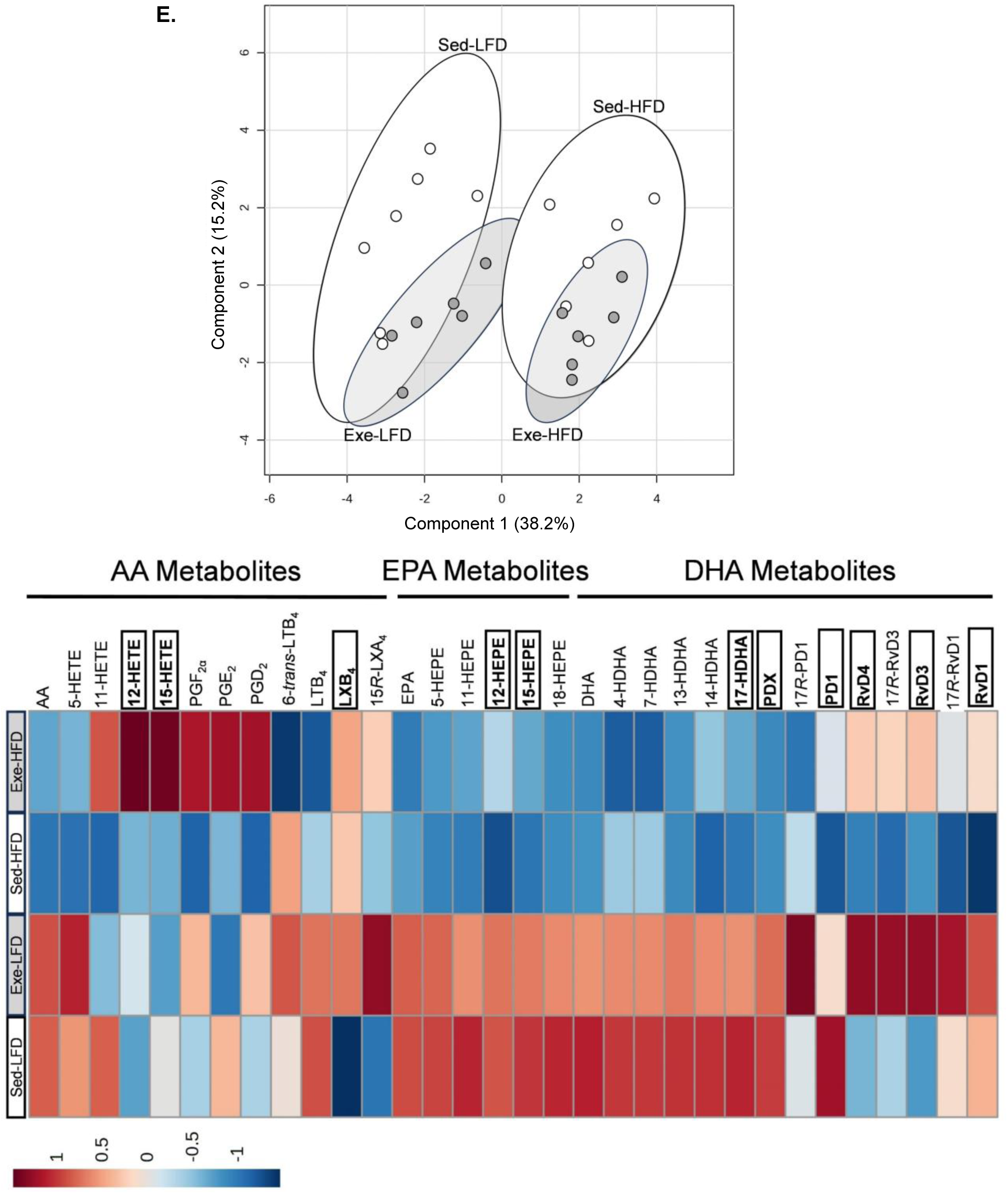
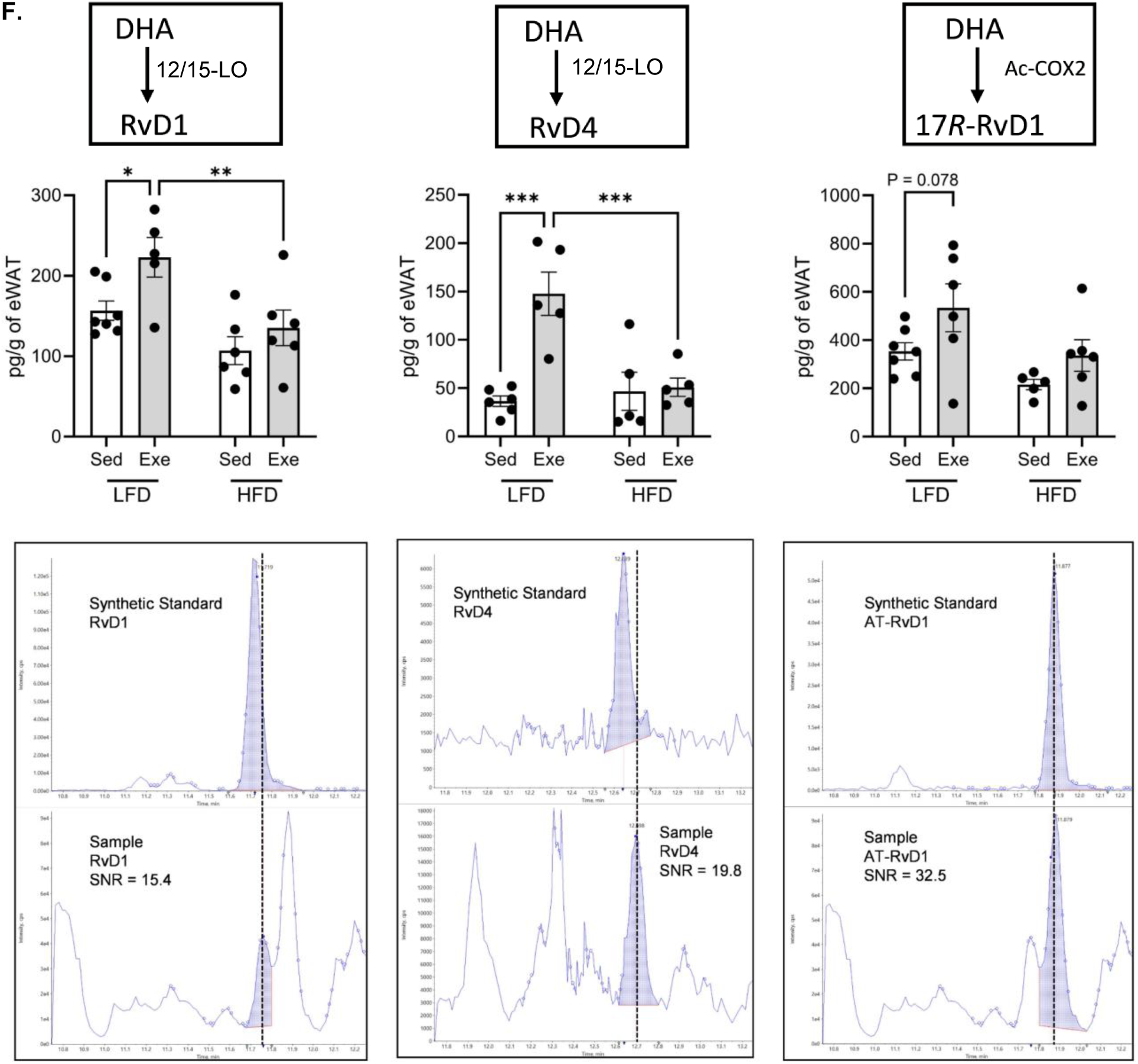
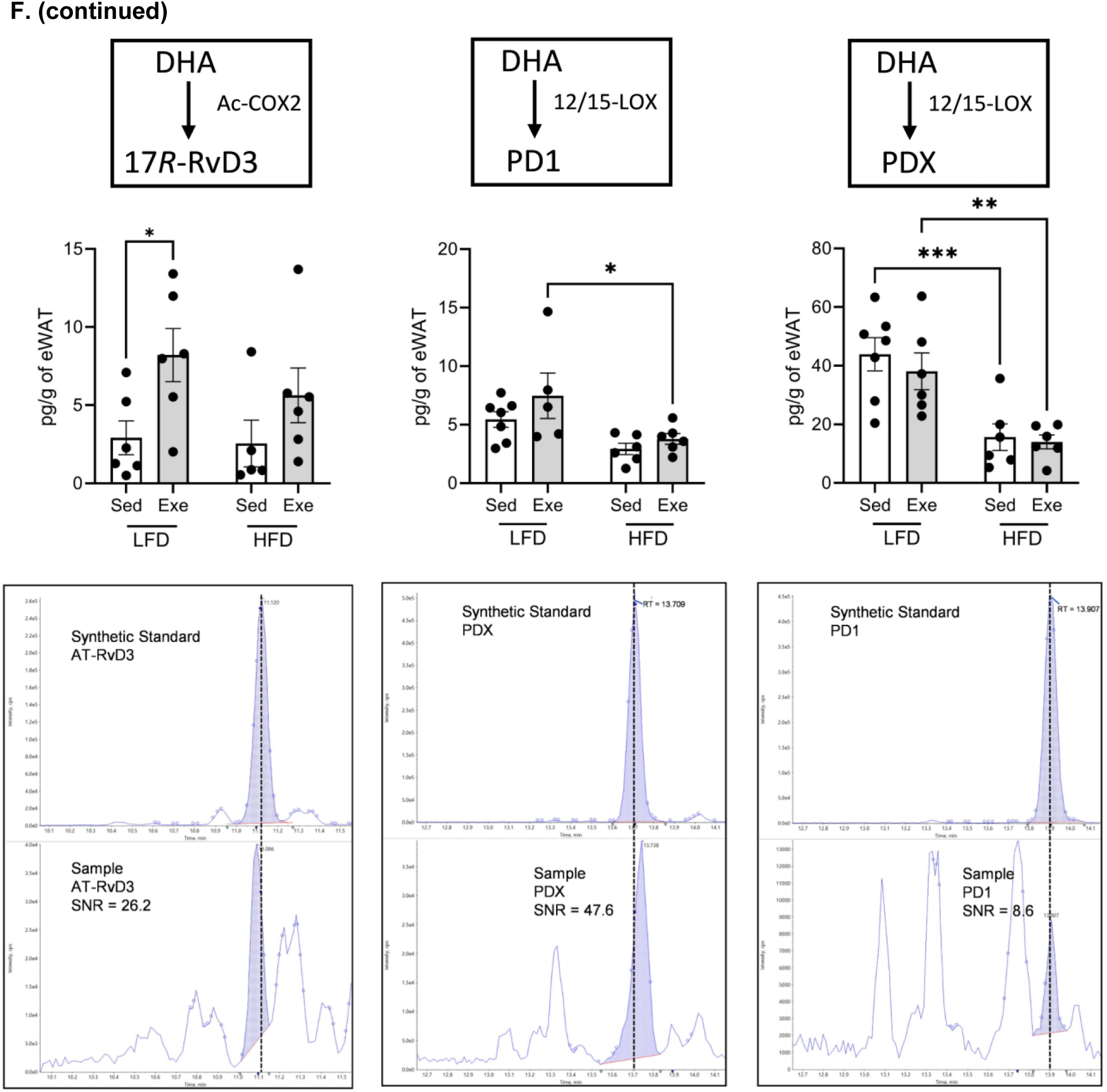
Exercise-stimulated *Alox15* expression, M2 macrophage content and SPM biosynthesis in adipose tissue is diet-dependent. (**A**) Body weights tracked throughout 6 weeks of either LFD or HFD feeding and concomitant 4 weeks of either sedentarism or exercise in male mice. (**B**) A graded, maximal exercise capacity test (ECT) was administered at baseline and after 4 weeks of exercise on the respective dietary group. Distance and work performed were measured at the end of the ECT, and blood lactate levels were measured before and after the ECT fatigue point in exercise-trained male mice fed either LFD or HFD. In sedentary male mice, blood lactate was only measured at baseline before the start of the ECT, as they did not undergo exercise capacity testing. (**C**) *Alox15* mRNA expression in whole eWAT and in F4/80^+^ ATM isolated from eWAT in the respective diet and exercise group. (**D**) %F4/80^+^ CD11c^+^ CD301^-^ (M1) and %F4/80^+^ CD11c^-^ CD301^+^ (M2) ATMs (as percent of all F4/80^+^ cells) in the respective diet and exercise group (**E**) 2D scores plot of sparse Partial Least Squares-Discriminant Analysis (sPLS-DA) and heat map of lipid mediator metabolipidomics from the eWAT from the respective diet and exercise group. (**F**) Illustration summarizing the biosynthetic pathway of the quantified D-series Resolvins and Protectins from their precursor molecule, DHA, and their respective quantifications below. Extracted ion chromatograms from a representative synthetic standard and eWAT sample showing the quantified peak areas, retention times and signal-to-noise ratios (SNR) are provided below each quantified analyte. Data expressed as mean ± SEM, *n* = 8-12 for all groups in (**A-B**); n = 4-8 for all groups in (**C**); n = 4-6 for all groups in (**D**); n = 6-7 in all groups in (**E**); n = 5-7 for all groups in (**F**). \**P*<0.05, \*\**P*<0.01, \*\*\**P*<0.001, \*\*\*\**P*<0.0001; One-way ANOVA with Holm-Šídák post-test in the last datapoint measured at week 4 (**A**), repeated measure one-way ANOVA (**B**), two-way ANOVA with Holm-Šídák post-test (**C, D, F**). sPLS-DA 2D scores plot and heatmap of group averages of analyte concentrations (**E**).

### 2.7 Urinary Epinephrine Measurements

Urinary epinephrine levels were measured following exercise training as previously described.^22^ Briefly, urine samples were collected following 4 weeks of exercise training and frozen at -80 °C immediately. Samples were then thawed on ice, vortexed, diluted with 0.2% formic acid, and mixed with deuterated epinephrine internal standard (±)-epinephrine-d_6._ Epinephrine was detected on an UPLC-MS/MS instrument (Acquity UPLC H-Class and Xevo TQ-S Micro Triple Quadrupole Mass Spectrometer with an electrospray ionization source; Waters Corporation, Milford, MA, USA). Epinephrine was separated from other sample constituents on an Acquity UPLC HSS PFP column (150 × 2.1 mm, 1.8 μm) (cat. no. 186005968, Waters Corporation) using a binary gradient composed of 0.2% formic acid (solvent A) and methanol (solvent B). Three MRM transitions were used – 1 for quantification, 1 for confirmation and 1 for labeled internal standard. These MRMs were scheduled around the retention time of the analyte and no less than 12 data points were collected per peak. Epinephrine was quantified using a ratiometric calibration curve method, using an 8-point calibration curve.

### 2.8 RT-qPCR

SVF and ATM extracts were aliquoted and centrifuged, and the cell pellets were resuspended in 300 μl RTL buffer containing β-mercaptoethanol (cat. no. 63689-25ML-F; Millipore-Sigma). For analysis of catecholamine biosynthetic enzymes, freshly dissected adrenal glands (1 gland per mouse) were homogenized in 350 µl RTL buffer containing β-mercaptoethanol. Whole-cell RNA was isolated and purified using RNeasy Mini Kit (cat. no. 74106, Qiagen; Hilden, Germany) following manufacturer’s instructions, performing DNA digestion with RNase-free DNase I Set (cat no. 79256, Qiagen) before eluting. RNA concentration and purity were determined by UV absorbance spectrophotometry using a NanoDrop 2000c (Thermo Fisher). The following amounts of RNA were used as template for cDNA synthesis for the indicated tissue and cell lysates: eWAT SVF, 27.72 ng; eWAT adipocyte fraction, 46.2 ng; eWAT ATM, 39.6 ng. cDNAs from adrenal gland, eWAT SVF, and adipocyte fractions were prepared by reverse transcription PCR using MultiScribe Reverse Transcriptase with the random primers scheme for initiation of cDNA synthesis (part of the High Capacity cDNA RT Kit; cat. no. 43-688-13, Thermo Fisher). cDNAs synthesis in ATM cell lysates was performed by reverse transcription PCR using SuperScript™ IV VILO™ Master Mix. Validated, commercially available PCR primers were purchased from Qiagen (RT^2^ qPCR Primers, cat. no. 330001) or from Thermo Fisher (TaqMan® Gene Expression Assays for *Alox15* in eWAT ATMs; cat. no. Mm00772337_m1 (4448484)) and used for quantitative PCR assays (see primer information in Table S5 of Supplemental Methods). Custom-made primers for *Hprt* were made by Integrated DNA Technologies and used to normalize relative mRNA expression of the indicated genes in adrenals, eWAT SVF, and adipocyte fractions. Primers for *Hprt* used to normalize relative *Alox15* mRNA expression in eWAT ATM were purchased from ThermoFisher (TaqMan® Gene Expression Assays; cat. no. Mm01324427_m1 (4331182)) (see primer information in Table S6 of Supplemental Methods). PowerUP SYBR Green Master Mix (cat. no. A25776, Thermo Fisher) with Dual-Lock Taq DNA polymerase and ROX as passive reference dye was used for quantitative RT-PCR of adrenal glands, eWAT SVF, and adipocytes. TaqMan^TM^ Fast Advanced Master Mix (cat. no. 4444557, Thermo Fisher) with AmpliTaq™ Fast DNA Polymerase, uracil-N-glycosylase, and ROX passive reference dye was used for quantitative RT-PCR of eWAT ATM. All reactions were run with manufacturer’s recommended acquisition settings on an Applied Biosystems QuantStudio 5 (Thermo Fisher). The 2 ^-ΔΔCT^ method was used to calculate relative expression following normalization to the housekeeping gene *Hprt*.

### 2.9 Immunoblotting analysis on adrenal glands

One whole adrenal gland was homogenized in 100 µl of lysis buffer composed of 1× RIPA buffer (cat. no. R0278-50ML, Millipore Sigma), 1× Halt Protease inhibitor cocktail (cat. no. 87785, Thermo Fisher) and 1× Halt Phosphatase inhibitor cocktail (cat. no. 78420, Thermo Fisher). Protein concentration was determined using Pierce BCA protein assay kit (cat. no. 23227, Thermo Fisher). 75 μg of protein per sample was mixed with 1× Laemmli sample buffer (cat. no. 1610747, BioRad, Hercules, CA, USA), resolved on an Anykd™ Criterion™ TGX™ Precast Midi Protein SDS-PAGE gel (cat. no. 5671124, Bio-Rad) and electrotransferred onto Immun-Blot PVDF membranes (150-160 μg/cm^2^) (cat. no. 1620177, BioRad). Membranes were blocked with 1% (w/v) non-fat milk (cat. no. M-0841, LabScientific, Danvers, MA, USA) in TBS-T buffer (TBS (cat. no. 1706435, BioRad) containing 0.1% v/v Tween-20 (cat. no. BP337-500, Fisher Scientific, Waltham, MA, USA) for 1h. The membrane was then probed overnight with mouse anti-PNMT primary antibody (1:1000 dilution in 2% w/V BSA in TBS-T; cat. no. NBP2-00688, Novus Biologicals, Centennial, CO, USA), or rabbit glucocorticoid receptor beta (GR-β)-A antibody (1:1000 dilution in 5% w/v BSA in TBS-T; cat. no. BS-13385r, ThermoFisher). After overnight incubation, the PNMT blot was washed 3 times in TBS-T and incubated with anti-mouse HRP-conjugated secondary IgG (1:1000 dilution; cat. no. 7076S, Cell Signaling Technology, Danvers, MA, USA) in 5% w/v non-fat milk for 1h. After overnight incubation, the GR-β blot was washed 3 times with TBS-T and incubated with anti-rabbit HRP-conjugated secondary IgG (1:2000 dilution; cat. no. HAF008, R&D Systems) in 5% w/v non-fat milk (cat. no. M-0841, Lab Scientific) for 1 h. The membrane was washed 3 times in TBS-T and then incubated with Pierce ECL Plus (cat. no. 32132, Thermo Fisher) and imaged using an MYECL Western Blotting Detection System (Thermo Fisher). Blot images and bands were quantified using ImageLab software (version 6.10, build 7; BioRad).

### 2.10 Histological analysis of adrenal glands

Freshly dissected adrenal glands were fixed in 10% formalin (cat. no. 23245684, Fisher Scientific) for 24 h, before transferring to 70% ethanol (EtOH) (cat. no. E7023-500ML, Sigma-Aldrich). Intact adrenals were processed in a Sakura Tissue Tek VIP 2000 (Sakura Finetek USA, Torrance, CA, USA) followed by paraffin embedding. Paraffin tissue blocks were deparaffinized by incubating 3 times (in separate containers), 5 min each, in Xylenes (cat. no. X5-1, Fisher Scientific), followed by 5 3-min EtOH incubations, with EtOH concentrations decreasing from 100% to 50% (one EtOH concentration per container/bath) in a gradient fashion. After a 3-min deionized water (diH_2_O) rinse, adrenals were stained with Hematoxylin (cat. no. 22-220-100, Fisher Scientific) for 3 min, rinsed with diH_2_O for 5 min, and stained with Bluing Reagent stain (cat. no. 22-220-106, Fisher Scientific) by quickly dipping adrenals 10 times – 1 s each time – and finally rinsed with diH_2_O for 5 min. Adrenals were then counter-stained with Eosin-Y (cat. no. 22-220-104, Fisher Scientific) for 40 sec before quickly dipping them 5 times in 95% EtOH, and then 10 times in 100% EtOH, in order to dehydrate tissues. Tissues were then cleared by incubating twice in xylene for 5 min each. Finally, adrenals were mounted on a cover slip and gross histological examination and imaging of tissue was performed on a Keyence BZ-X810 imaging system (Keyence Corp., Osaka, Japan) at 20× and 40× magnification using Brightfield mode.

### 2.11 Generation of mouse transformed BMMs and M2(IL-4) and M1(LPS/INF-γ) polarization for flow cytometric analysis

Immortalized bone-marrow-derived mouse macrophage (BMM) cell lines were established by injecting the bone marrow of WT C57BL/6J mice with the mouse recombinant J2 retrovirus containing the v-myc and v-raf oncogenes, as previously described. ^25^ Cells were cultured in R5 media composed of the following: L-Gln-free RPMI640 (cat. no. 21870076, Thermo Fisher) supplemented with 5% FBS, 1× Glutamax (Thermo Fisher cat. no. 35050061) and 1% v/v HEPES (from 1M stock, cat. no. H0887-100ML, Sigma-Aldrich), in a humidified incubator at 37°C and 5% CO_2_. BMM were maintained at a low seeding density of 1E6 cells per T-75 flask in 10 mL R5 media and allowed to reach ∼75% confluency before passaging. Cell stocks were cryopreserved at -80°C in R5 media with 10% v/v DMSO at 1E6 cells per mL in 2 mL cryogenic vials (cat. no. 430659, Corning, Corning, NY, USA) using Nalgene Cryo 1 °C, ‘Mr. Frosty’ Freezing Containers (cat. no. 5100-0036, Thermo Fisher).

The BMM were polarized in vitro to corroborate their ability to upregulate canonical M1 (CD11c) and M2 (CD301) markers in response to the respective polarization stimuli (LPS/INF-γ for M1 and IL-4 for M2), and to investigate their response to SPMs. For M1(LPS/INF-γ) polarization experiments, BMM were subcultured at passage 6-15, at a density of 0.65E6 cells per 12-well plate well in 1 mL media, and were incubated with 1 nM RvD1 (cat. no. 10012554, Cayman Chemical Company) for 1 h prior to, and throughout a 24-h incubation with 5 ng/mL LPS (cat. no. L2880-10MG, Sigma-Aldrich) and 12 ng/mL INF-γ (cat. no. 315-05, Pepro Tech, Cranbury, NJ, USA). Upon RvD1 and LPS/INF-γ incubation, cells were harvested for flow cytometric analysis (see Fig. S7). For M2(IL-4) polarization experiments, BMM were subcultured at passage 6–15, at a density of 0.5E6 cells per 12-well plate well in 1 mL media, and incubated with 5 ng/mL IL-4 (cat. no. 404-ML-010, R&D Systems, Minneapolis, MN, USA) (or vehicle control) and either 166.6 μM BSA-conjugated palmitate (stock of 5 mM palmitate:0.8 mM BSA at a 6:1 molar ratio; cat. no. 29558, Cayman Chemical) or 27.8 µM BSA control (cat. no. A7030-50G, Sigma-Aldrich) for 24 h. This BSA control concentration was chosen to match the BSA concentration in the BSA-palmitate conjugate solution (6:1 palmitate:BSA molar ratio). After 24 h, without washing away IL-4, BSA-Palmitate, or respective vehicle, 25 µM etomoxir (cat. no. E1905-5MG, Sigma-Aldrich) (or vehicle) was incubated for 30 min before the addition of 1 nM RvD1. BMM were incubated for an additional 24 h before collection for immunophenotyping via flow cytometry. Briefly, culture media was aspirated, cells washed with 1 mL Ca^2+^- and Mg^2+^- free DPBS, and 400 µL of accutase (cat. no. 07922, STEMCELL Technologies) per well was incubated for 5 minutes at 37 °C before scraping off cells, washing and resuspending in FACS buffer for flow cytometric analysis, as previously described (see Methods subsection “Flow Cytometric Analysis of eWAT SVF or isolated ATM”). Note that, in addition to the antibodies previously listed, CD45-FITC (at a working concentration of 20 µg/ml) was added to this flow cytometric panel.

### 2.12 Mouse elicited peritoneal macrophage (PM) extraction and Seahorse XF Assay

PM cells were isolated as previously described.^23^ Upon isolation, 1E5 cells were seeded per well (XFe96 plate) in 180 µL for 2 h in a nutrient- and serum-limited growth media composed of Seahorse basal DMEM (cat. no. A1443001, Agilent, Santa Clara, CA, USA) supplemented with 0.5 mM D-glucose (555 mM stock; Sigma-Aldrich cat. no. G8644-100ML), 1 mM L-Gln (200 mM stock; cat. no. 103579-100, Agilent), 0.5 mM L-carnitine (cat. no. C0283-1G, Sigma-Aldrich), 1% FBS and 1% v/v penicillin/streptomycin (from a mixed stock containing 10,000 U penicillin and 10 mg/ml streptomycin; cat. no. P4333-20ML, Sigma-Aldrich). After the 2 h incubation, the cells were washed 2× with 200 µL of assay media composed of Seahorse basal DMEM containing 2 mM D-glucose and 0.5 mM L-carnitine followed by addition of medium containing 1 nM RvD1, 1 nM RvD4 (cat. no. 13835, Cayman Chemical Company) or vehicle. The cells were then incubated for 1 h at 37°C and at normal atmospheric CO_2_ partial pressure in a humidified incubator. After SPM incubation, cells were incubated with either 166 µM BSA-conjugated palmitate, 166 µM BSA-conjugated oleate or 27.8 µM BSA control for 30 min before the start of a mitochondrial stress assay. For this assay, OCR and ECAR measurements were taken continuously in 6-min intervals, with the first 3 min spent mixing and the last 3 min spent reading. Basal measurements were taken for 59 min (to allow for the respective fatty acids to be taken up and catabolized), and then, to probe mitochondrial respiratory parameters, oligomycin, FCCP, and Antimycin-A/Rotenone solutions were successively added (without washing away the previous compound) to each well at the following respective time points (relative to the start of the 1^st^ basal reading) – 60 min, 78 min, 98 min. The following working concentrations of mitochondrial inhibitors were used, as determined by dose response titrations performed beforehand in separate experiments (data not shown): 1.5 µM oligomycin, 1.5 µM FCCP, 10 μM Antimycin-A, 1 µM Rotenone (MilliporeSigma cat. no. O4876-5MG, C2920-10MG, A8674-50MG and R8875-1G, respectively). This assay was performed on an Agilent Seahorse XFe96 extracellular flux analyzer. The following formulas were used to calculate the indicated mitochondrial respiratory parameters: Non-Mitochondrial OCR = minimum OCR value achieved after Antimycin-A + Rotenone (AA/R) injection; Basal OCR = (last OCR value before oligomycin injection) – (Non-Mitochondrial OCR); Maximal OCR = (max OCR value after FCCP injection) – (Non-Mitochondrial OCR); ATP-Linked OCR = (last OCR value before oligomycin injection) – (minimum OCR value achieved after oligomycin injection); Spare Respiratory Capacity = (Max OCR) – (Basal OCR); Spare Respiratory Capacity as % of Basal = [(Max OCR) / (Basal OCR)] × 100%; H^+^ Leak = (minimum OCR value after oligomycin injection) – (Non-Mitochondrial OCR); % Coupling Efficiency = [(ATP-Linked OCR) / (Basal OCR)] x 100%.

### 2.13 Seahorse XF assay on isolated SVF cells

SVF cells were isolated as described above in Methods subsection 2.4. Due to the per-mouse cell yield being lower than required for all the groups and technical replicates, cells from 2 animals were combined into a single biological replicate sample. SVF cells in post-digestion buffer were washed twice with pre-warmed XF assay media composed of basal Seahorse DMEM supplemented with 10 mM D-glucose, 2 mM L-Gln and 1 mM sodium pyruvate. 1.5E6 Cells were seeded per well (XFe96 plate) with 1 nM RvD1 or vehicle control in 180 µL and incubated for 2h at 37°C and at normal atmospheric CO_2_ partial pressure. 166.6 µM BSA-conjugated palmitate or 27.8 µM BSA control were added 30 min before the start of the Seahorse XF assay. The mitochondrial stress assay was carried out as described in this manuscript, with the exception that basal OCR and ECAR was measured for 19 min, and thus oligomycin, FCCP and antimycin-A/Rotenone solutions were added at the following times (relative to the start of the 1^st^ basal reading) – 20 min, 39 min, 59 min. The following working concentrations of mitochondrial inhibitors were used, as determined by dose response titrations performed beforehand in separate experiments (data not shown): 3 µM oligomycin, 2 µM FCCP, 10 µM Antimycin-A, 1 µM Rotenone.

### 2.14 Statistical Analysis

Results are expressed mean ± standard error of the mean (SEM) of n observations, where n represents the number of biological replicates per experimental group. All animals were randomly assigned to experimental groups and age matched with proper controls. All statistical comparisons were performed with GraphPad Prism v9.3.1 (GraphPad Software, La Jolla, CA, USA). Statistical comparisons between 2 groups were conducted using either paired or unpaired two-tailed Student’s *t* test, while comparisons of multiple groups were performed using one-way ANOVA or two-way ANOVA, where appropriate, with Holm-Šídák’s post hoc tests, as indicated. Normality and data homoscedasticity were assumed. In all cases, statistical significance was set at *P* < 0.05. Sample sizes and *p* values (for the indicated pair-wise comparisons) for the respective graphs are indicated in figure legends. Statistical outliers were determined using Grubbs’ test with alpha = 0.05.

## 3. RESULTS

### 3.1 Exercise stimulates duration dependent changes in the expression of 12/15-lipoxygenase and anti-inflammatory macrophage content in white adipose tissue

Because adipose tissue is critical for whole-body metabolism and that exercise exerts dynamic and pleiotropic effects, we first examined duration dependent changes of exercise on white adipose tissue. For this, mice were randomly assigned to either sedentarism (Sed control group) or to 1, 2, or 4 weeks of aerobic treadmill exercise. Following training, administration of a graded maximal exercise capacity test demonstrated that 1, 2, and 4 weeks of training resulted in an exercise-duration dependent improvement in distance ran and work performed when compared with sedentary control animals (Fig. 1A). Improvements in exercise capacity were associated with exercise duration dependent changes in eWAT mass, heart mass, liver mass, kidney mass, and spleen mass (Fig. S1A), despite no changes in total body weight (Fig. 1B).

We next investigated the effects of exercise on lipid mediator production in adipose tissue via targeted lipidomics on eWAT. Following data dimensionality reduction analysis, we observed that concentrations of lipid mediators in adipose tissue after 4 weeks of exercise training clustered independently from the sedentary group and the 1 wk and 2 wk exercise groups (Fig. 1C). This differential clustering was driven, in part, by multiple 12/15-LO-derived lipid mediators (Fig. 1C). We next questioned whether *Alox15* (murine 12/15-lipoxygenase) expression was altered by exercise training. To this end, we observed enhanced *Alox15* mRNA expression in the adipose tissue stromal vascular fraction (SVF) following 4 wk of exercise training with a commensurate decrease in *Alox15* expression in adipocytes (Fig. 1D). Because *Alox15* is dynamically expressed in macrophages depending on polarization, we questioned whether adipose tissue macrophages (ATMs) were responsible for the exercise-enhanced *Alox15* expression we observed in the SVF fraction. To test this, we isolated F4/80^+^ ATMs from non-ATM F4/80^-^ SVF cells from the 4 wk exercise and sedentary groups and observed increased expression of *Alox15* mRNA specifically in the F4/80^+^ ATM fraction (Fig. 1F).

Given that *Alox15* expression is upregulated in anti-inflammatory macrophages,^26,27^ we wondered whether the exercise-induced *Alox15* expression observed in ATMs was associated with an exercise-induced shift in ATM polarization and abundance. Interestingly, 4 wk of exercise training resulted in a shift in the balance of M2 and M1 macrophages with increased abundance of anti-inflammatory (F4/80^+^CD301^+^CD11c^-^) ATMs and decreased abundance of inflammatory (F4/80^+^CD301^-^CD11c^+^) ATMs, despite having no significant effect on total macrophage (F4/80^+^) abundance (Fig. 1G). Collectively, these data demonstrate a dynamic shift in the expression of *Alox15* and abundance of CD301^+^ anti-inflammatory macrophages in adipose tissue following 4 wk of aerobic exercise training.

### 3.2 Exercise-stimulated Alox15 expression, M2 macrophage content and SPM biosynthesis in adipose tissue is altered by diet

We previously showed that exercise-stimulated resolvin D1 biosynthesis and enhanced macrophage phagocytosis enhances resolution of acute inflammation.^22^ Additionally, our group and others have shown that long duration HFD consumption in mice leads to a failure in pro-resolving pathways are associated with obesity-induced adipose tissue inflammation.^9,10,12,13,28,29^ Therefore, we investigated whether a shorter duration of high fat feeding (6 wk) influences exercise-stimulated *Alox15* expression in ATMs. As shown in Fig. 2A, HFD feeding did not significantly alter body composition, whereas exercise increased lean mass with a concomitant decrease in fat mass in both dietary groups (Fig. S2A). Importantly, diet had no effect on exercise-enhanced performance as shown by a similar increase in distance, work performed, and increase in blood lactate in a graded, maximal exercise capacity test administered before and after 4 weeks of training (Fig. 2B). Using this model of short duration HFD feeding, we observed no alterations in fasting blood glucose levels (Fig. S2B), GTT, or ITT (Fig. S2C), suggesting no significant changes in systemic glucose or insulin sensitivity. Despite no changes in glucose tolerance or insulin sensitivity, it is interesting to note that in LFD-fed mice, exercise training decreased non-fasting insulin levels, an effect that was abrogated by HFD feeding (Fig. S2D). Additionally, adiponectin, an important insulin-sensitizing hormone whose release is stimulated by physical activity (in both humans^30–32^ and mice^33–35^), particularly in obese individuals,^30,36,37^ was significantly downregulated in HFD-fed sedentary mice relative to their LFD-fed counterparts, and exercise was unable to rescue defective adiponectin secretion (Fig. S2D). These changes in insulin and adiponectin secretion may be indicative of early HFD-induced perturbations before the onset of overt adipose tissue inflammation and systemic insulin resistance.

Given the importance of AT in maintaining whole-body energy homeostasis, and its susceptibility to HFD-induced perturbations, we investigated exercise- and diet-induced changes in AT. Consistent with data presented in Figure 1, we found that 4 wk of exercise increases *Alox15* expression in both eWAT and isolated F4/80+ ATMs of control diet (LFD)-fed mice, an effect abrogated by HFD feeding (Fig. 2C). Moreover, LFD fed animals subjected to exercise demonstrated higher abundance of anti-inflammatory M2-polarized (F4/80^+^CD301^+^CD11c^-^) ATMs. This exercise-induced effect was abrogated by HFD feeding (Fig. 2D). Interestingly, the inhibition of exercise-induced anti-inflammatory CD301+ macrophages in the adipose tissue by 6 wk of HFD feeding occurred in the absence of overt adipose tissue inflammation more commonly observed in longer-duration HFD feeding,^38^ as evidenced by the lack of an increase in the abundance of pro-inflammatory M1(F4/80^+^ CD301^-^ CD11c^+^) ATMs (Fig. 2D).

We next questioned whether exercise-induced upregulation of *Alox15* mRNA expression in LFD fed animals resulted in changes in pro-resolving 12/15-LO-derived lipid mediator levels in adipose tissue. Using targeted LC-MS/MS, we quantified levels of eicosanoid- and docosanoid-derived lipid mediators. After performing a dimensionality reduction analysis, we observed that exercise in LFD fed animals resulted in modest separation, which was inhibited in HFD fed animals (Fig. 2E). We observed that HFD feeding broadly inhibited docosahexaenoic acid (DHA) and eicosapentaenoic acid (EPA) products, many of which are 12/15-LO-derived (Figs. 2E, S2E). Additionally, multiple arachidonic acid (ArA)-derived metabolites were upregulated in exercised animals fed a HFD when compared with LFD fed animals (Fig. 2E). Moreover, levels of multiple 12/15-LO derived SPMs, such as RvD1, RvD4, 17*R*- RvD1, and 17*R*-RvD3, were elevated by exercise in LFD animals compared with sedentary controls, which was inhibited by HFD feeding (Figs. 2F). Furthermore, HFD feeding inhibited the biosynthesis of the SPM PD1 and its isomer PDx, independent of exercise intervention (Fig 2F). Collectively, these data suggest that HFD consumption inhibits exercise-stimulated *Alox15* expression, SPM production, and CD301+ anti-inflammatory macrophage abundance in adipose tissue.

### 3.3 12/15-LO derived SPMs stimulate fatty acid β-oxidation that is directly coupled with CD301 expression in macrophages

We next sought to understand the molecular mechanisms by which exercise-stimulated SPM biosynthesis increases the abundance of CD301+ anti-inflammatory M2 adipose tissue macrophages. We have recently reported that exercise-stimulated SPM biosynthesis enhances macrophage mitochondrial respiration – consistent with an M2 metabolic phenotype.^23,39^ Given this, we questioned whether SPMs, fatty acid β-oxidation, and CD301 expression were coupled in macrophages. To test this, we induced M2 macrophages using IL-4 in CD45+F4/80+ bone marrow-derived macrophages (BMMs) and assessed protein expression of CD301 (Fig. 3A). Interleukin 4 (IL-4) induced CD301 expression, which was further potentiated by the addition of palmitate to the culture media (Fig. 3A). The addition of 25 μM of the carnitine palmitoyltransferase 1 (CPT-1) (rate limiting step in β oxidation) inhibitor Etomoxir (see Fig. S6) reversed palmitate-induced CD301 expression (Fig. 3A). These results suggest that protein expression of CD301 is directly coupled with fatty acid β oxidation in macrophages.

**Figure 3.**
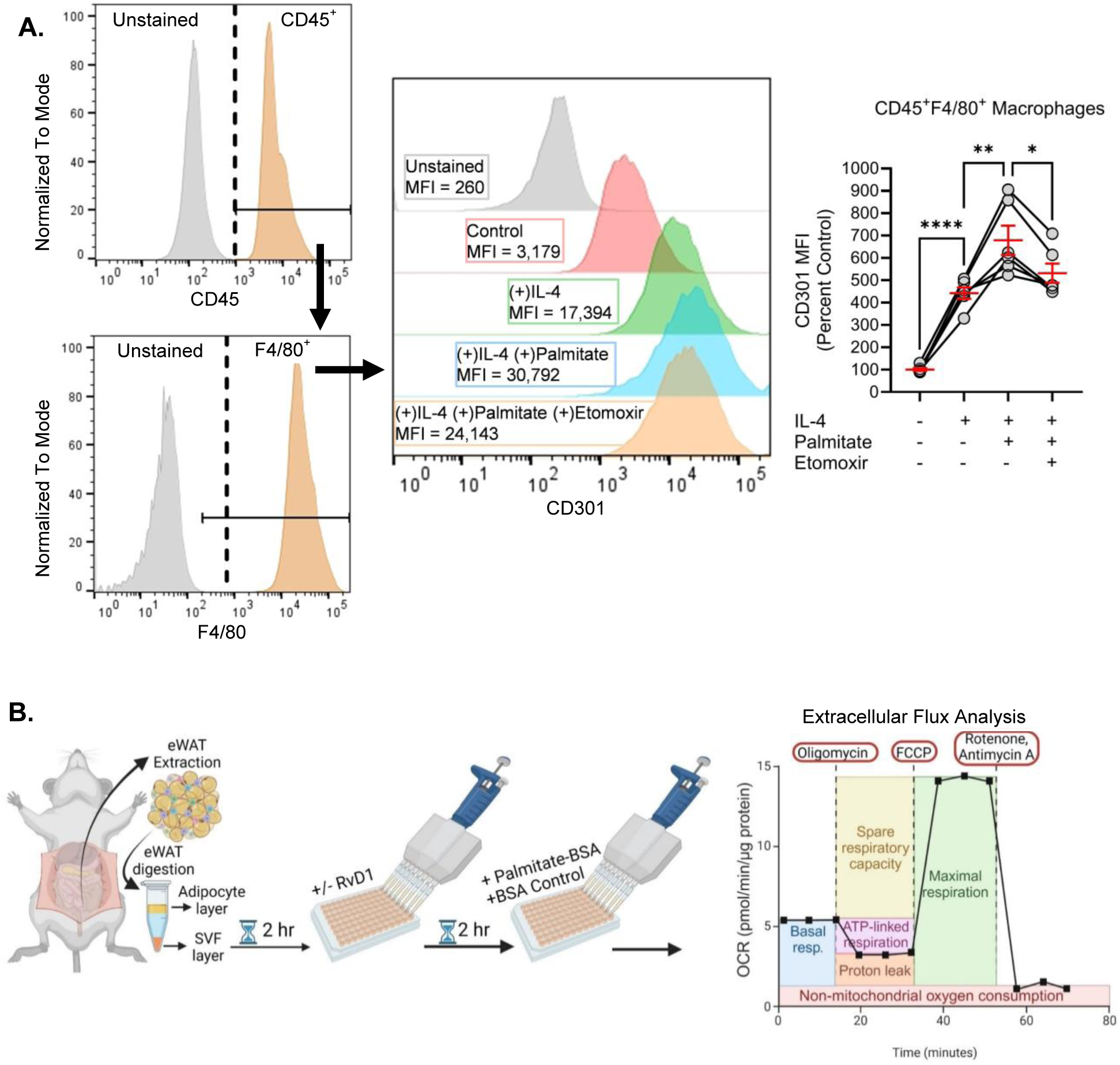
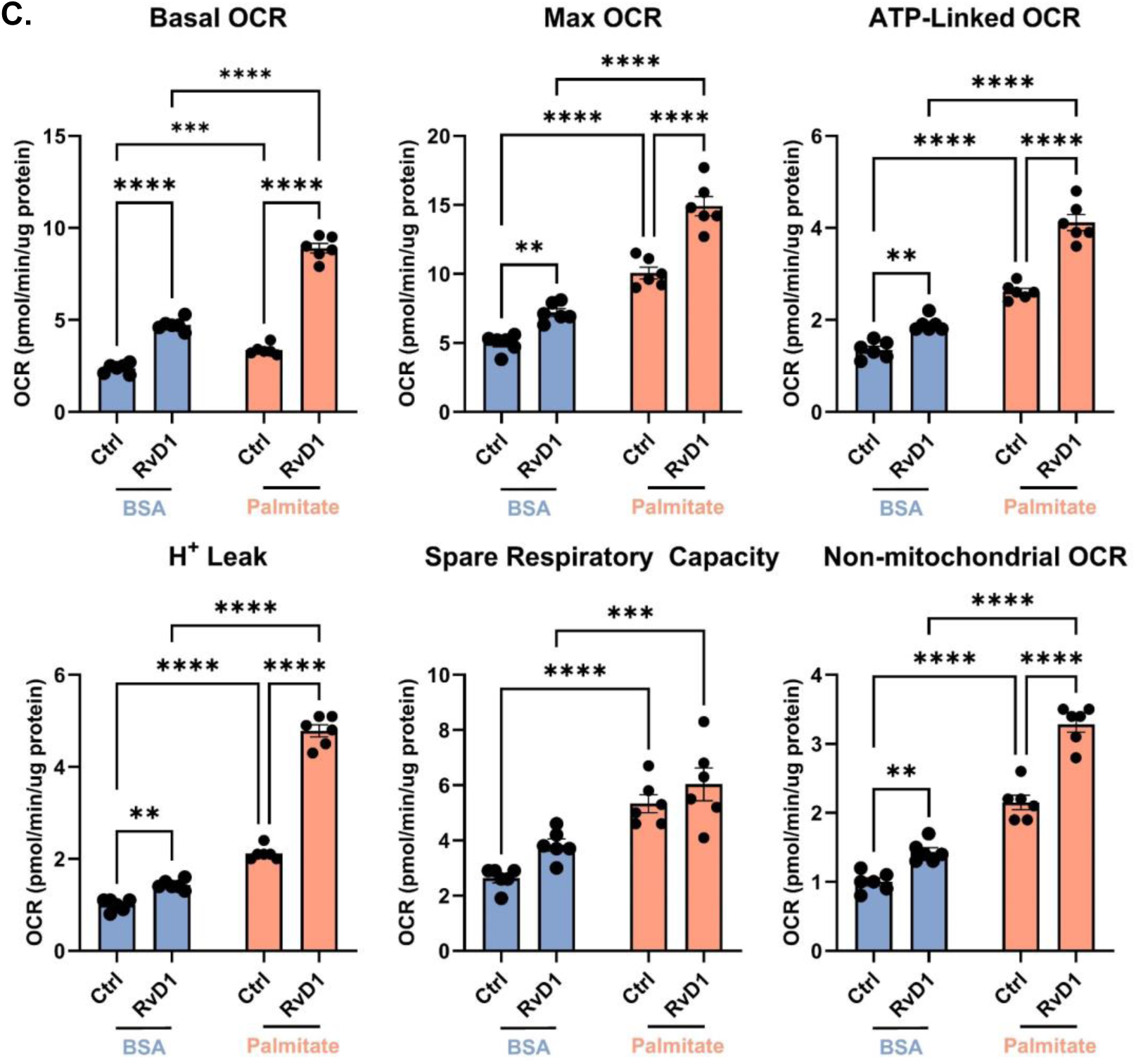
12/15-LO-derived SPMs stimulate fatty acid β-oxidation, which is directly coupled with CD301 expression in macrophages. (**A**) Gating strategy used to identify CD45^+^ F4/80^+^ CD301^+^ M2(IL-4) macrophages and subsequently measure CD301 expression levels via mean fluorescence intensity (MFI) of the identified population under the indicated treatments. (**B**) Experimental scheme outlining the isolation of eWAT SVF cells from naïve male mice, subsequent +/- RvD1 and +/-palmitate treatments, and subsequent measurements of mitochondrial respiratory parameters using a mitochondrial stress assay – namely (**C**) Basal OCR, maximal OCR, ATP-linked OCR, H^+^ leak, spare respiratory capacity, and non-mitochondrial respiration. Data expressed as mean ± SEM; *n* = 6 (**A**), n = 6 (**C**); \**P*<0.05, \*\**P*<0.01, \*\*\**P*<0.001, \*\*\*\**P*<0.0001; One-way ANOVA with Holm-Šídák post-test (**A**), two-way ANOVA with Holm-Šídák post-test (**C**).

We next questioned whether the 12/15-LO derived SPM, RvD1, promotes β oxidation in SVF cells isolated from adipose tissue (Fig. 3B). Indeed, we found that incubation with RvD1 stimulated basal OCR, maximal OCR, ATP-linked OCR, H+ leak, and non-mitochondrial OCR (Fig. 3C). Importantly, oxygen consumption was further potentiated by the presence of palmitate (Fig. 3C), indicating the RvD1 stimulates β oxidation of the fatty acid palmitate. We also tested this phenomenon in elicited macrophages (Fig. S3) and found that basal oxygen consumption rate was enhanced with RvD1 and RvD4 (both SPMs were found to be elevated in adipose tissue with exercise in LFD fed animals (Fig. 2F-2G)), which was further potentiated in the presence of palmitate or oleate (Fig. S3B), two of the most abundant fatty acids. Collectively, these results suggest that SPMs stimulate β oxidation, which is directly coupled with expression of CD301 in macrophages.

### 3.4 HFD inhibits PNMT expression and promotes adrenal fatigue

We have previously shown that exercise-induced catecholamine release and α1-AR-dependent signaling on peritoneal macrophages is necessary and sufficient to mediate the exercise-induced upregulation of *Alox15* and RvD1 in macrophages.^22^ Thus, we questioned whether the HFD-induced inhibition of exercise-induced *Alox15* expression and SPM biosynthesis in ATMs is caused by alterations in catecholamine biosynthesis. Although no gross histological abnormalities were observed in adrenal glands of HFD-fed relative to LFD-fed animals (Fig. 4A), we found that HFD significantly reduced *Pnmt* mRNA expression, a key biosynthetic enzyme that produces epinephrine from norepinephrine (Fig. 4B, 4C). We next sought to confirm downregulation of protein expression and performed immunoblotting of PNMT in adrenal glands isolated from LFD and HFD fed animals subjected to exercise or sedentary conditions. To this end, we found that HFD feeding resulted in downregulation of PNMT at the protein level which was not reversed by exercise training compared with LFD fed controls (Fig. 4D). Additionally, HFD feeding inhibited the exercise-induced enhancement of epinephrine levels (Fig. 4E).

**Figure 4.**
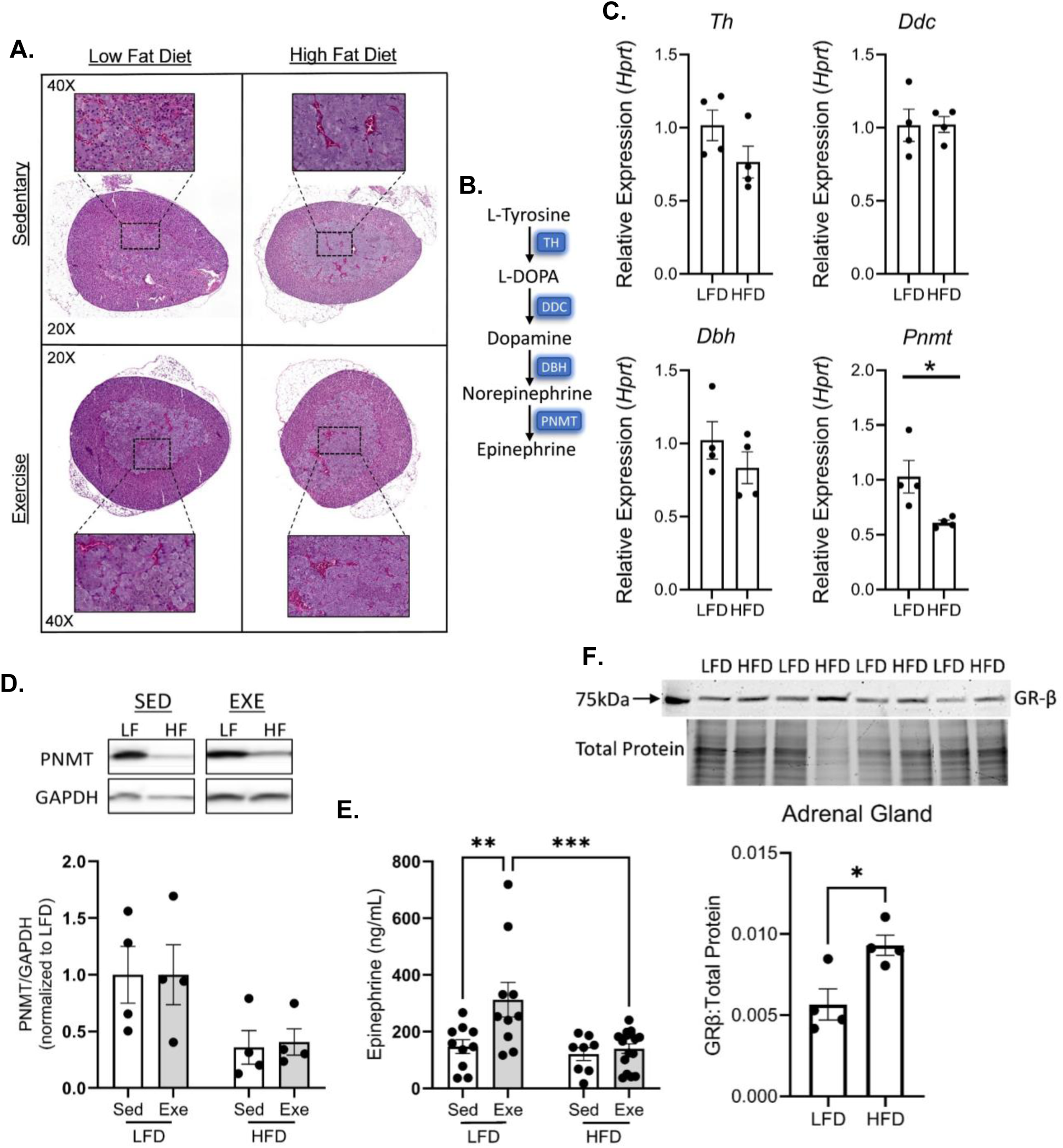
**HFD promotes adrenal fatigue and inhibits PNMT expression**. (**A**) H&E staining of adrenal glands from LFD-Sed and HFD-Sed male mice. (**B**) Graphical outline of the biosynthetic pathway for epinephrine. (**C**) mRNA expression of adrenal biosynthetic enzymes *Th*, *Ddc*, *Dbh* and *Pntm*, relative to *Hprt*, in either LFD- or HFD-fed sedentary male mice. (**D**) Ratio of PNMT to GAPDH protein expression in adrenal glands (with representative immunoblots) for either LFD- or HFD-fed, sedentary or exercised male mice. PNMT/GAPDH ratios are normalized to the LFD-Sed group. (**E**) Urinary epinephrine levels in either LFD- or HFD-fed, sedentary or exercised male mice. (**F**) Western immunoblotting quantification and representative blots of adrenal GR-β. Data expressed as mean ± SEM; *n* = 4 (C-D, F), n = 8-16 (E); \**P*<0.05, \*\**P*<0.01, \*\*\**P*<0.001; two-tailed Student’s *t*-test (B, F), two-way ANOVA with Holm-Šídák post-test (C-D).

To explore possible molecular mechanisms by which HFD induced the observed suppression in catecholamine release, we measured the mRNA expression of 3 transcription factors TFAP2A, EGR-1, and glucocorticoid receptor NR3C1 – known to regulate PNMT transcription.^40–44^ As shown in Fig. S4A, short duration HFD feeding did not significantly alter mRNA expression of these transcription factors relative to LFD-fed counterparts. Interestingly, however, HFD feeding increased glucocorticoid receptor-β (GR-β) expression (Fig. 4F), an inactive glucocorticoid receptor splice variant that is known to inhibit GR-α signaling^45–48^ and inhibit PNMT expression and epinephrine biosynthesis. Further investigation of this signaling pathway is required to establish a possible causal link between HFD feeding and the observed adrenergic insufficiency.

### 3.5 Epinephrine-induced increase in CD301+ M2 ATM content in HFD-fed animals is dependent on 12/15-LO

Next, we questioned if supplementing HFD fed animals with epinephrine could rescue HFD-induced adrenergic insufficiency and restore a pro-resolving state in the AT, and whether this is dependent on 12/15-LO. To test this, mice were fed HFD for 5 wk and treated for 1 week (while kept on HFD) with daily doses of either vehicle, epinephrine alone, or pre-treated with a molecular inhibitor of 12/15-lipoxygenase – ML351 – 2 h before epinephrine treatment (Fig. 5A). The dose and duration of ML351 treatment were chosen to achieve peak tissue concentrations, based on published pharmacokinetics and pharmacodynamics studies.^49,50^ We found several 12/15-LO-derived metabolites of ArA (15-HETE and 12-HETE) (Fig S5A), DHA (14-HDHA, MaR1, 17-HDHA) (Fig. S5B) and EPA (12-HEPE and 15-HEPE) (Fig. S5C) were significantly lower in the eWAT of epinephrine- and ML351-treated mice relative to mice treated with epinephrine alone. Interestingly, we found that epinephrine treatment stimulated F4/80^+^ ATM recruitment, with increases in both M1 (F4/80^+^ CD11c^+^ CD301^-^) and M2 (F4/80^+^ CD11c^-^ CD301^+^) abundance (Fig. 5B). To test for sex differences, we subjected female mice to the same HFD and epinephrine treatments and observed a similar increase in the total F4/80^+^ ATMs with an increase in only M1 (F4/80+CD11c+CD301-) and not M2 (F4/80^+^ CD11c^-^ CD301^+^) macrophages (Fig S5E). Given the lack of M2 macrophage induction in females, we focused the remainder of our studies on male mice. In agreement with the flow cytometry data, epinephrine treatment elevated the mRNA expression of known M2 polarization markers – namely *Arg-1*,^51,52^ *Adgre-1*,^53^ *Mgl-1*,^52^ and *Ym-1*,^52,54,55^ within the SVF (Fig. S5D). Highlighting the role of 12/15-LO in mediating exercise-induced CD301+ macrophage abundance in the AT, blocking 12/15-LO-derived lipid mediator production with ML351 before epinephrine administration inhibited CD301+ M2 macrophages abundance, while increasing M1 pro-inflammatory ATM content. Collectively, these data suggest that epinephrine is necessary to promote macrophage recruitment to adipose tissue; however, 12/15-LO is required for promoting CD301+ M2 macrophage abundance.

**Figure 5.**
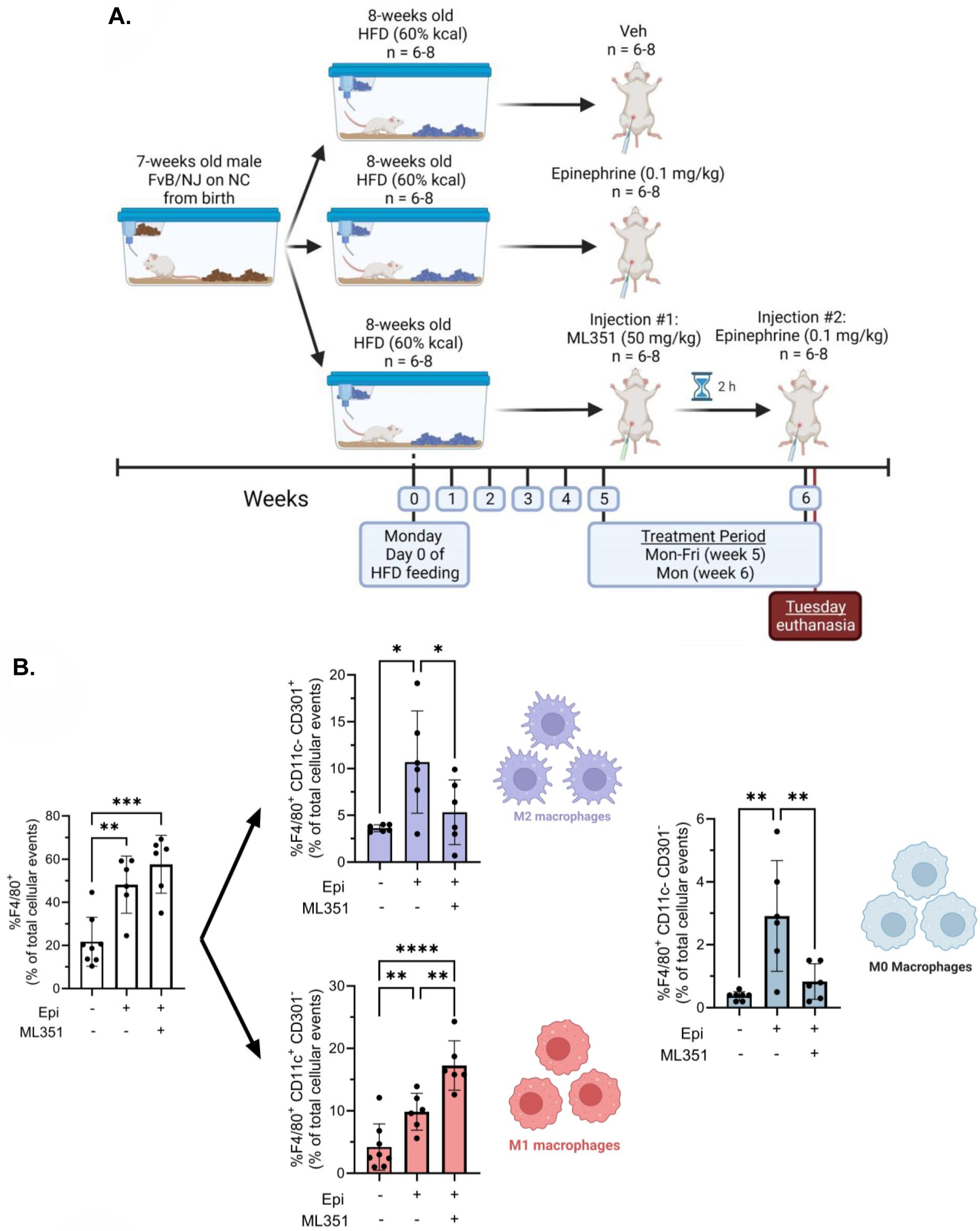
Epinephrine-induced increase in M2 ATM content in HFD-fed animals is dependent on 12/15-LO activity. (**A**) Experimental scheme outlining time points for the administration of diets as well as epinephrine and ML351 treatments in male mice. (**B**) %F4/80^+^ ATMs, %F4/80^+^ CD11c^+^ CD301^-^ (M1 ATMs), %F4/80^+^ CD11c^-^ CD301^+^ (M2 ATMs) and %F4/80^+^ CD11c^-^ CD301^-^ (M0 ATMs), quantified as percent of total cellular events within the eWAT SVF compartment of male mice, via flow cytometry. Data expressed as mean ± SEM; *n* = 6-8; \**P*<0.05, \*\**P*<0.01, \*\*\**P*<0.001; One-way ANOVA with Holm-Šídák post-test.

## 4. DISCUSSION

In this study, we find that exercise training increases adipose tissue macrophage *Alox15* expression, D-series resolving production, and the abundance of CD301+ M2 macrophages. Each of these exercise-induced benefits in adipose tissue were abrogated when animals concomitantly consumed HFD. Additionally, HFD feeding inhibited PNMT expression in adrenal glands and exercise-stimulated epinephrine, which when added back to HFD-fed mice resulted in increased total macrophage abundance with increases in both F4/80+CD301+CD11c-M2 and F4/80+CD301-CD11c+ M1 macrophages. Inhibition of 12/15-LO by ML351 resulted in elevated epinephrine-induced M1 macrophages and decreased M2 macrophages. Mechanistically, we also report that macrophage CD301 expression is directly coupled with fatty acid β oxidation, which is stimulated by 12/15-LO derived SPMs. Collectively, these findings suggest that exercise stimulates epinephrine production, which enhances macrophage 12/15-LO expression and SPM production to promote fatty acid oxidation and CD301+ macrophage abundance in adipose tissue.

Among immune cell types that populate the adipose tissue stromal vascular fraction, macrophages play a pivotal role in maintaining homeostasis during physiological (i.e. stress- and fasting-induced lipolysis, cold exposure) and pathological conditions. These crucial tasks include the control of pre-adipocyte differentiation, maintenance of vascular integrity, extracellular matrix remodeling, modulating angiogenic responses during hypertrophy and hyperplasia, clearance of apoptotic adipocytes, reuptake and temporary storage (or “buffering”) of adipocyte-derived FFAs during lipolysis, among others.^56–58^ ATM, whether tissue-resident or monocyte-derived, are dynamically programmed by the ever-changing AT microenvironmental milieu and are transcriptionally and phenotypically rewired to respond to such changes.

Recent advancements in the field of macrophage biology have challenged the existing dogma that macrophages exist in either one of two phenotypic extremes, i.e. classically activated, proinflammatory (M1) vs. alternatively activated, anti-inflammatory (M2) macrophages. Rather, they exist as a complex phenotypic spectrum that, in the example of ATMs, encompasses many highly specialized subsets that fulfil critical roles to meet dynamic tissue needs, such as during nutrient storage versus release.^56,58,59^ For example, during periods of fasting-induced lipolysis, tissue-resident vascular-associated ATMs are initially depleted followed by chemoattraction of circulating monocytes that differentiate and display higher phagocytic potential. These newly recruited, highly phagocytic ATMs play a critical role in lysosomal-mediated lipid uptake and removal of dead adipocytes.^60–63^ During prolonged periods of weight loss, however, reductions in central adiposity are reproducibly accompanied by a decrease in circulating inflammatory markers as well as a restoration of M2-like ATM abundance and an overall amelioration of the inflammatory tone in adipose tissue.^64–66^ Likewise, during periods of cold stress-induced thermogenesis, adaptive visceral white AT remodeling occurs, during which time a dynamic M1-to-M2 phenotypic switch occurs. This highly coordinated response is dependent on CSF1 receptor signaling and adrenergic stimulation^67^ and contributes to the dynamic changes in ATMs in response to changing environmental cues in adipose tissue.

In the present study we find that exercise-stimulated catecholamine release, pro-resolving lipid mediator biosynthesis, and increased abundance of M2(F4/80^+^CD301^+^CD11c^-^) ATMs is abrogated by high fat diet-induced adrenergic insufficiency. Consistent with the published findings that fasting and physical activity stimulate β-AR-dependent lipolysis that in turn induces ATM recruitment to “buffer” released FFA,^63,68^ we observed an elevation in F4/80^+^CD301^+^CD11c^-^ M2 ATMs with exercise-stimulated catecholamine release. Even though some reports have shown that exercise exerts its anti-inflammatory effects via epinephrine-stimulated downregulation of inflammatory cytokine release in innate immune cells (i.e. TNF and IL-1β),^69^ to our knowledge, this is the first report to show a direct involvement of exercise-stimulated epinephrine stimulating pro-resolving lipid mediator production and a concomitant enrichment in the F4/80^+^CD301^+^CD11c^-^ M2 ATM population in visceral adipose tissue. These findings constitute a novel mechanism by which exercise mediates protective benefits.

We find that short-term HFD feeding leads to dysregulated adrenergic signaling in response to exercise with reduced exercise-stimulated epinephrine release, AT SPM biosynthesis, and reduced presence of M2 (F4/80^+^CD301^+^CD11c^-^) anti-inflammatory macrophages. Moreover, it is well established that HFD-induced obesity causes disruption to the hypothalamic-pituitary-adrenal (HPA) axis, thus affecting the production of several hormones from the adrenal cortex.^70,71^ For example, increased adiposity is associated with HPA axis overactivation, evidenced by heightened ACTH and cortisol release in response to exogenous stimulation.^72–75^ In contrast, several reports investigating acute responses to exercise demonstrate that obese individuals have blunted epinephrine release and diminished adrenergic-stimulated lipolysis in visceral fat depots.^76–82^ Despite this associative evidence in humans, the molecular mechanisms undergirding impaired adrenergic signaling in obesity and their effect on adipose tissue immune cells remain incompletely understood. It’s interesting to speculate that catecholamine resistance in adipose tissue with concomitant elevations in epinephrine release may promote adrenal medulla fatigue and decreased epinephrine release as we observed with HFD feeding. Concomitant with a decrease in epinephrine release, a loss in catecholamine signaling in the AT may also result from HFD-induced enrichment of sympathetic neuron-associated macrophages (SAM). These tissue-resident ATMs are known to express the noradrenaline transporter SLC6A2 and the catecholamine catabolizing enzyme MAO-A,^56,68,83^ thus impairing β-adrenergic-stimulated lipolysis. Findings presented here however suggest that dysregulated adrenal medulla function with HFD feeding inhibits exercise-stimulated epinephrine release.

In the present study, we chose to quantify M2 macrophages based on their expression of the Galactose-type C-type lectin, CLEC10A (CD301), a marker of M2 macrophages in both humans and mice, which is involved in immune cell interactions and pathogen recognition.^84,85^ Although C-type mannose receptor 1 (CD206), also termed MRC1, is a more commonly used M2 marker to identify macrophages in humans and mice,^84^ recent evidence suggests that both CD206^+^ and CD206^+^CD11c^+^ double positive ATM populations produce pro-inflammatory cytokines such as IL-6 and IL-8.^56,86,87^ Additionally, CD206+ M2 macrophages do not express a diagnostic M2-like gene signature profile relative to the characteristic M1 gene signature profile of macrophages expressing the diagnostic M1 marker CD11c.^88^ This recent evidence brings into question the use of CD206 as a pan marker of M2 macrophage polarization.

The relationship between cellular metabolism and macrophage immunophenotype has been well described. For example, M2 macrophages primarily rely on mitochondrial respiration for ATP generation, whereas M1 macrophages primarily derive ATP via aerobic glycolysis. We find that mitochondrial β-oxidation of palmitate and oleate, two of the most abundant FFAs in the AT microenvironment,^89^ is potentiated by treatment with 12/15-LO derived SPMs – consistent with an M2 phenotype.^39^ Furthermore, addition of palmitate to M2(IL-4)-polarized macrophages *in vitro* further enhanced the expression of CD301, which was abrogated upon fatty acid oxidation inhibition with etomoxir. These exercise-stimulated, and SPM-mediated changes, in ATM bioenergetics (previously reported^23^) – consistent with an M2 phenotype constitute a possible novel mechanism by which ATMs are able to carry out FFA buffering and removal of dying adipocytes, as occurs during caloric restriction or under exercise-induced adrenergic stimulation.^63,89^ That 12/15-LO derived SPMs may be critical to maintain adipose tissue homeostasis is further supported by the findings of Kwon et al., who demonstrated in a model of adrenergic-stimulated lipolysis that macrophages from *Alox15*-null animals display a failure to clear dying adipocytes and maintain tissue homeostasis as a result of downregulation in genes associated with lipid uptake and metabolism.^90^

### 4.1 Conclusion

In summary, our findings suggest that the adrenergic stimulated pro-resolving effects of exercise in adipose tissue are modified by diet. Specifically, we find that six weeks of HFD feeding is sufficient to inhibit exercise-induced catecholamine release, in-turn inhibiting exercise stimulated 12/15-LO-derived SPM biosynthesis and decreasing the abundance of pro-resolving M2 ATMs. This feeding duration is too short to promote overt obesity, adipose tissue inflammation, or insulin resistance, which is typically observed after 12-24 weeks or more of HFD feeding.^91–93^ This suggests that HFD feeding disrupts the normally present inflammation resolution programs that help maintain the energy storage and neuroendocrine function of the adipose tissue.^8–11,13^

## NON-STANDARD ABBREVIATIONS AND ACRONYMS

12/15-LO: 12/15-lipoxygenase
AA/R: antimycin-A and rotenone mix
ACTH: adrenocorticotropic hormone
AR: adrenergic receptor
ArA: arachidonic acid
AT: adipose tissue
ATM: adipose tissue macrophage
BMM: bone marrow-derived macrophages
BSA: bovine serum albumin
CPT-1: palmitoyltransferase-1
DEXA: dual-energy X-ray absorptiometry
DHA: docosahexaenoic acid
DMSO: dimethyl sulfoxide
ECAR: extracellular acidification rate
ECT: exercise capacity test
EDTA: Ethylenediaminetetraacetic acid
EPA: eicosapentaenoic acid
EPI: enhanced product ion
ESB: EasySep^TM^ buffer
EtOH: ethanol
eWAT: epididymal white adipose tissue
FACS: fluorescence activated cell sorting
FBS: fetal bovine serum
FCCP: carbonyl cyanide-p-trifluoromethoxyphenylhydrazone
FFA: free fatty acid
GAPDH: Glyceraldehyde 3-phosphate dehydrogenase
GR-β: glucocorticoid receptor β
GTT: glucose tolerance test
HFD: high-fat diet
HPA: hypothalamic-pituitary-adrenal axis
HPLC: high-pressure liquid chromatography
HPRT: Hypoxanthine-guanine phosphoribosyltransferase
I.P.: intraperitoneal
IL-10: interleukin-10
IL-1Ra: interleukin-1 receptor antagonist
IL-1β: interleukin-1β
IL-4: interleukin-4
IL-6: interleukin-6
INF-γ: interferon-γ
IS: internal standard
ITT: insulin tolerance test
LC-MS/MS: liquid chromatography tandem mass spectrometry
LFD: low fat diet
LLOD: lower limit of detection
LLOQ: lower limit of quantification
LM: lipid mediator
LPS: lipopolysaccharide
LXA_4_: lipoxin A_4_
MAO-A: monoamine oxidase-A
MaR1: maresin
MCP-1: macrophage chemoattractant protein 1
MeOH: methanol
MFI: mean fluorescence intensity
MRM: multiple reaction monitoring
NC: normal chow
OCR: oxygen consumption rate
PD1: protectin D1
PDx: protectin Dx
PLSDA: partial least-square determinant
PM: peritoneal macrophage
PNMT: phenylethanolamine N-methyltransferase
PUFA: polyunsaturated fatty acid
RBP4: retinol binding protein 4
RCF: relative centrifugal force
RT: retention time
RvD1: resolvin D1
RvD4: resolvin D4
SAM: sympathetic neuron-associated macrophages
Sed: sedentary
SEM: standard error of the mean
SNR: signal-to-noise ratio
sPLS-DA: sparse partial least square – discriminant analysis
SPMs: specialized pro-resolving lipid mediators
SVF: stromal vascular fraction
T2D: type-2 diabetes
TNF-α: tumor necrosis factor alpha
UPLC-MS/MS: ultra high-pressure liquid chromatography tandem mass spectrometry
WAT: white adipose tissue
Wk: week
XF: extracellular flux

## ACKNOWLEDGEMENTS

The authors thank the University of Louisville, Center for Cardiometabolic Science Flow Cytometry Core. Additionally, the authors would like to thank the animal and administrative support staff for their tireless efforts in aiding the progress of science. Illustrated components of figures 1, 3, 5, S3, and S5, along with the graphical abstract, were created using BioRender.com.

## SOURCES OF FUNDING

This work was supported in part by National Institutes of Health grants GM127495 (J.H.), GM127495-S1 (J.H.), HL130174 (B.G.H.), HL106173 (M.S.), and ES028268 (B.G.H.). The Center for Cardiometabolic Science is supported by the National Institutes of Health grant GM127607. Ernesto Pena Calderin is the recipient of an National Research Service Award Ruth L. Kirschstein F31 Fellowship (DK131920).

EPC, JH, JZ, NB, WL, BS and MS conducted experiments and analyzed data. EPC and JH wrote the manuscript with input from all the authors. BGH assisted on experimental protocol design and data analysis/interpretation. JH conceived, designed, and supervised the research and writing of the manuscript.

## SUPPLEMENTAL MATERIALS

Supplemental Methods

Tables S1-S6

Figures S1-S7

## HIGHLIGHTS

- Exercise-stimulated specialized proresolving lipid mediator biosynthesis (SPM) and proresolving M2 macrophage content in adipose tissue is diet-dependent.
- 12/15-LO-derived SPMs stimulate fatty acid β-oxidation, which is directly coupled with CD301 expression in ATMs
- HFD promotes adrenal fatigue and inhibits PNMT expression
- Epinephrine-induced increase in M2 ATM content in HFD-fed animals is dependent on 12/15-LO activity

